# A Heat-Shock Inducible System for Flexible Gene Expression in Cereals

**DOI:** 10.1101/2020.05.29.123554

**Authors:** Sophie A. Harrington, Anna E. Backhaus, Samantha Fox, Christian Rogers, Philippa Borrill, Cristobal Uauy, Annis Richardson

**Affiliations:** John Innes Centre, Norwich Research Park, Norwich NR4 7UH, United Kingdom; ENSA, Sainsbury Laboratory, University of Cambridge, Cambridge CB2 1LR, United Kingdom; School of Biosciences, University of Birmingham, Birmingham B15 2TT, United Kingdom; Institute of Molecular Plant Sciences, University of Edinburgh, Edinburgh EH9 3BF, United Kingdom

**Keywords:** Inducible promoters, Cre recombinase, transgenic, heat shock, inducible expression, clonal sectors, wheat, barley, cereals, grasses

## Abstract

**Background:** Functional characterisation of genes using transgenic methods is increasingly common in cereal crops. Yet standard methods of gene over-expression can lead to undesirable developmental phenotypes, or even embryo lethality, due to ectopic gene expression. Inducible expression systems allow the study of such genes by preventing their expression until treatment with the specific inducer. When combined with the Cre-Lox recombination system, inducible promoters can be used to initiate constitutive expression of a gene of interest. Yet while these systems are well established in dicot model plants, like *Arabidopsis thaliana*, they have not yet been implemented in grasses.

**Results:** Here we present an irreversible heat-shock inducible system developed using Golden Gate-compatible components which utilises Cre recombinase to drive constitutive gene expression in barley and wheat. We show that a heat shock treatment of 38 °C is sufficient to activate the construct and drive expression of the gene of interest. Modulating the duration of heat shock controls the density of induced cells. Short durations of heat shock cause activation of the construct in isolated single cells, while longer durations lead to global construct activation. The system can be successfully activated in multiple tissues and at multiple developmental stages and shows no activation at standard growth temperatures (~ 20 °C).

**Conclusions:** This system provides an adaptable framework for use in gene functional characterisation in cereal crops. The developed vectors can be easily adapted for specific genes of interest within the Golden Gate cloning system. By using an environmental signal to induce activation of the construct, the system avoids pitfalls associated with consistent and complete application of chemical inducers. As with any inducible system, care must be taken to ensure that the expected construct activation has indeed taken place.

## Background

Historically, transformation in cereal crops such as wheat and barley has been hampered by low efficiencies and high costs (1). Yet recent advances in transformation techniques have shown remarkable improvements of transformation efficiency (1). As a result, functional characterisation of cereal genes increasingly uses transgenic approaches. Commonly, genes are characterised using standard over-expression systems, where the gene of interest is placed under the control of a constitutive promoter (2). In some cases, however, ectopic over-expression can be deleterious or even embryo lethal, especially when dealing with genes involved in development.

A common approach to overcome such difficulties involves the use of an inducible system, whereby the gene of interest is placed under the control of an inducible promoter. The gene is thus only expressed upon application of the required inducer. These systems are useful when dealing with genes whose overexpression may be deleterious for the plant as well as to induce gene expression in specific tissues or at specific developmental times (3). Chemicals such as β-estradiol (4, 5), ethanol (3, 6), and glucocorticoids such as dexamethasone (7, 8) have been used successfully to drive inducible transgenic systems in plants. Environmentally-driven systems have also been established, utilising promoters which are induced under specific abiotic stresses, such as heat (9, 10), cold (11), or drought (12).

While most of these inducible systems have been implemented in *Arabidopsis*, within the cereals environmentally-induced promoters have been most well characterised. A barley heat shock promoter, *HvHSP17*, was used to successfully drive the heat-shock induced expression of the reporter gene GUS in wheat (10). The promoters of two cold-responsive genes, *OsWRKY71* and *TdCor39*, were used to drive cold-induced expression of the wheat *DREB3* gene in both barley and rice (11). Temperature-independent systems have also been established successfully, including the use of the barley drought-responsive promoter *HvDhn4s* (12).

Yet while these inducible systems have been successful in causing activation of a given gene of interest following application of the required stimulus, they are unable to cause constitutive, non-reversible activation of that gene. These are inherently transitory systems. As a result, these systems must be combined with additional components to allow for constitutive, inducible expression of a gene of interest. One of the most well characterised systems that allows for constitutive activation of a gene of interest is the Cre-Lox system. Initially developed for use in mice (13), but now regularly used in many species including plants (14, 15), Cre recombinase is an enzyme derived from the bacteriophage P1 which recombines DNA between two specific DNA sequences, named the *loxP* sites. This system can therefore be used to excise a desired segment of DNA and has been used in conjunction with inducible systems in model plants to drive the controlled, constitutive activation of gene expression following induction (16, 17). The benefits of the Cre-Lox system have been amply demonstrated in model species such as *Arabidopsis* (18–20) but have rarely been applied to cereal crops (16). We therefore aimed to develop a framework for a transgenic system in cereals that would allow the irreversible induction of a gene of interest at the desired developmental stages.

Here we have integrated a barley heat-shock specific promoter, *HvHSP17*, with the Cre-Lox system and shown that we can successfully activate the construct following heat shock treatment. Gene activation can be achieved in multiple tissues and distinct developmental stages, demonstrating that the system is versatile. Variable lengths of applied heat shock can control the density of activated cells, with short periods inducing single-cell construct activation and longer periods inducing global activation. Construct activation can be optimised for specific experimental setups to ensure that the desired level of activation occurs.

## Results

### Design of the construct

To develop a system that would allow for the irreversible induction of a given gene of interest at a specific time, we developed a Cre-Lox system under the control of an inducible promoter. Our aim was to have a construct that consisted of an inducible promoter that would drive expression of Cre recombinase upon application of the appropriate stimulus. Initially, and in the absence of the stimulus, only the reporter gene would be expressed under the control of a constitutive promoter (Figure 1A). Expression of Cre recombinase after the appropriate treatment (Figure 1B) then leads to the irreversible excision of the reporter gene, located on the same construct and flanked by two *loxP* sites (Figure 1C). Removal of the reporter gene then brings the gene of interest in proximity to a constitutive promoter that drives its expression in an irreversible manner (Figure 1D). Previously, the barley heat shock promoter *HvHSP17* has been shown to be induced upon treatment for two hours at 38 °C in wheat, with minimal background expression at lower temperatures (10). We therefore selected this promoter as a strong, consistent inducible promoter that could be used to drive expression of the Cre recombinase. We then developed two parallel versions of this construct, one for use in barley for use in microscopy (Figure 1E), and one for use in wheat to test the impact of premature *NAM-B1* expression on wheat development (Figure 1F).

**Figure 1:**
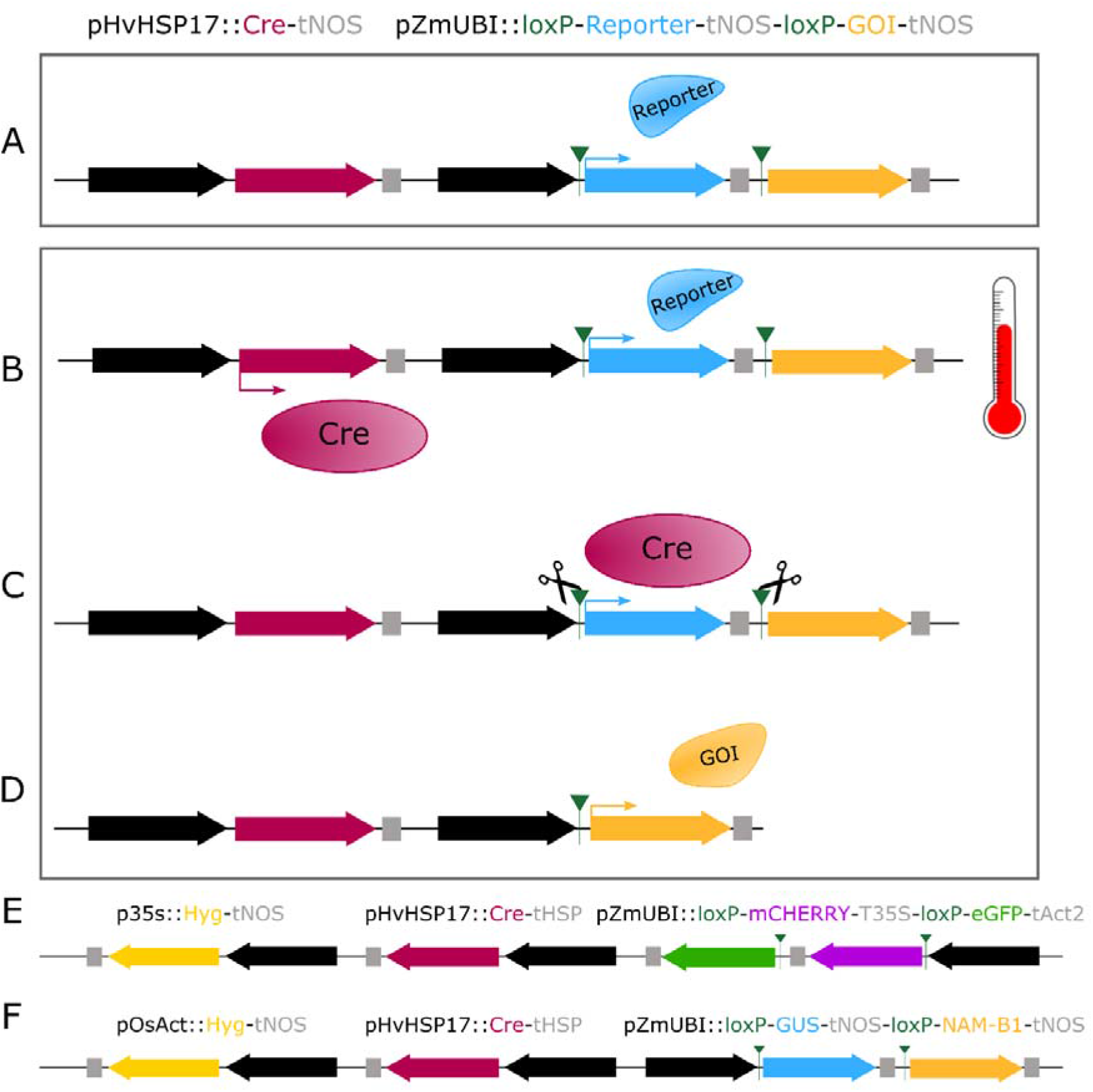
Schematic of the inducible Cre-Lox system. (A) The initial construct contains Cre recombinase (red), under the control of a stress inducible promoter, such as HvHSP17, and a constitutive expression promoter, here the maize Ubiquitin promoter (*pZmUbi*, black), driving expression of a reporter gene (blue) followed by an appropriate terminator (grey). The reporter gene is flanked by two *loxP* recombination sites (green triangles) and is followed by a second gene of interest (GOI; yellow) which is not expressed in the initial construct. (B) Following application of stress, in this case a heat shock treatment, expression of the *Cre* recombinase gene is induced. (C) The Cre recombinase protein carries out recombination at the *loxP* sites flanking the reporter gene. (D) Following excision of the reporter gene, the gene of interest is now in frame with the promoter sequence, and expression of the gene of interest is now under the control of the constitutive promoter. (E) A heat shock inducible construct, HS_GFP, utilised the barley *HvHSP17* promoter to drive expression of Cre recombinase following heat shock, leading to irreversible excision of the reporter gene, *mCHERRY*, and induction of *eGFP*. (F) This system has also been established in wheat, using *GUS* as the reporter gene which when excised leads to expression of *NAM-B1*. The *loxP* cassette is inverted in the wheat construct, relative to its position in the barley construct.

To track the activation of the construct at the cellular level, we required the use of fluorescent reporter proteins that could be effectively visualised using confocal microscopy without being obscured by the autofluorescence in mature barley tissues. To select the best fluorescent proteins for use in barley, we carried out a spectral scan on barley leaf and floral tissue to check for endogenous auto-florescence, which would interfere with fluorescent protein signals. Barley leaf and inflorescence tissue at different developmental stages was harvested and imaged using confocal laser microscopy to assess native fluorescence when excited with different wavelengths appropriate to CyPET (21), eGFP (22), and mCHERRY (23). For all three wavelengths, auto-fluorescence was only observed at high gain levels, except in the mature leaf tissue which had higher auto-fluorescence (Supplementary Figure 1). This suggested that CyPET, eGFP and mCHERRY were suitable to use in barley transgenics, especially when observing young tissues. However, the higher level of endogenous auto-fluorescence will need to be considered when imaging mature leaf tissue.

From these reporters, we selected *mCHERRY* for use as the reporter gene, flanked by the *loxP* sites, and *eGFP* for use as the induced gene, located downstream of *mCHERRY*. These genes were placed under the control of the constitutive maize Ubiquitin promoter, *ZmUbi*. In both cases, ER-targeting sequences were cloned at the N- and C-terminals of the fluorescent reporter genes to limit cell-cell movement of the fluorescent proteins. This construct was then cloned in tandem with the *Cre* gene containing the *U5* intron, driven by *HvHSP17,* and a selection cassette consisting of *Hygromycin* under the control of the constitutive 35S promoter (Figure 1E).

Alongside the barley construct, we developed a secondary version of the construct for transformation into wheat, which utilised a different reporter/gene system. As GUS had been successfully used as a reporter in previous tests of the heat shock promoter (10), we combined the *GUS* gene, flanked by the *loxP* sites, with the wheat NAC transcription factor *NAM-B1* (24). The remainder of the construct follows that of the barley construct (Figure 1F). Details of the specific components used to clone the barley and wheat constructs are presented in the methods section (Figure 8, Supplementary Figures 2 and 3).

### Characterisation of the heat shock construct

#### Transformation and copy number validation

T_0_ transformants of the wheat and barley constructs were obtained in the Fielder and Golden Promise cultivars, respectively, and screened for copy number. We identified 17 independent events which contained the wheat construct, HS_NAM-B1, and eight independent events which contained the barley construct, HS_GFP (Supplementary Table 1). Fourteen independent wheat lines and four barley lines were taken forward for copy number analysis. From these lines, four wheat lines with varied copy number were taken forward for the majority of further analysis at the T_1_ generation. One single-copy barley line was taken forward for analysis, from which only the homozygous lines containing a single insertion, and thus two copies, at the T_1_ generation were used for further study.

#### Heat shock induces activation of the Cre-Lox construct

To characterise the heat shock-inducible construct, we first tested whether the application of heat shock to positive transformants would be sufficient to induce expression of *Cre* recombinase. Based on previous work with the *HvHSP17* promoter (Freeman et al. 2011), we expected that two hours at 38 °C would be able to induce expression of *Cre*. Using T_1_ plants descended from the wheat 4-copy line 2020-2-1, we found that within two hours of heat shock treatment, *Cre* expression could already be detected (Supplementary Figure 4).

We investigated whether the expression of *Cre* was sufficient to cause successful excision of the reporter gene. In wheat seedlings before heat shock treatment, PCR of extracted genomic DNA using primers flanking the *loxP* sites (Figure 2A) amplified the expected 2490 bp region encompassing the reporter gene (Figure 2B). Following two hours of heat shock at 38 °C, a shorter band of 151 bp was observed (Figure 2A, B). This indicated that the Cre recombinase can excise the reporter gene between the two *loxP* sites. Of the 14 independent wheat lines tested with this assay, all those which expressed the *GUS* reporter gene also demonstrated successful construct excision following seedling heat shock (data not shown).

**Figure 2:**
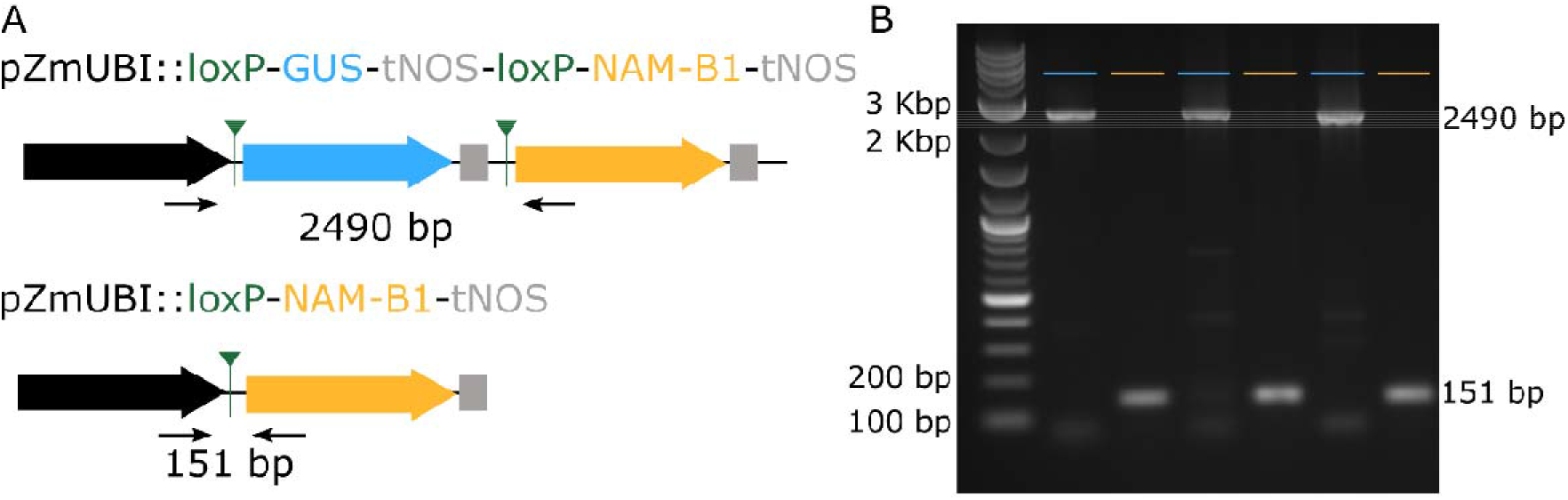
Heat shock induces excision of the reporter gene. (A) Excision of the *GUS* gene and expression of *NAM-B1* occurs following heat-shock and expression of *Cre* recombinase. The location of the primers used to validate the *GUS* excision are shown in black, alongside the expected band size. (B) Gel image shows the resultant DNA bands before (blue) and after (yellow) heat shock. This representative figure depicts the results from three independent lines, 2020-3-1, 2020-24-01, and 2020-21-01, at the T_1_ generation.

#### Heat shock leads to the expression of the gene of interest

As heat shock treatment leads to the successful excision of the reporter gene, we next investigated the expression levels of the gene of interest following recombination. We selected three independent transgenic lines containing the *NAM-B1* construct for further testing: 2020-20-02 (one copy), 2020-2-1 (four copies), and 2020-5-1 (ten copies). We also included the null copy control, 2010-54-01. Three or four individual T_1_ plants of each line were treated at the two-leaf stage either with two hours at 38 °C (heat shock; “HS”) or two hours at 20 °C (non-heat shock; “NHS”), and were sampled for RNA extraction two weeks after the treatment. There was no significant expression of *NAM-B1* in the null copy control under any condition (Figure 3A), as expected. While there was a statistically significant increase in expression in one of the positive transgenic lines before heat shock (“Pre”) compared to the 0 copy control (4 copy line; Student’s T-test, p < 0.05), the spread of gene expression observed was within the biological background levels observed in the 0 copy line. This suggests that no substantial gene expression is observed before heat shock application. However, after heat shock (“Post”) all three lines showed a significant (2020-2-1, 2020-5-1, p < 0.05) or near-significant (2020-20-02, p ~ 0.09) increase in *NAM-B1* above the baseline expression before treatment.

**Figure 3:**
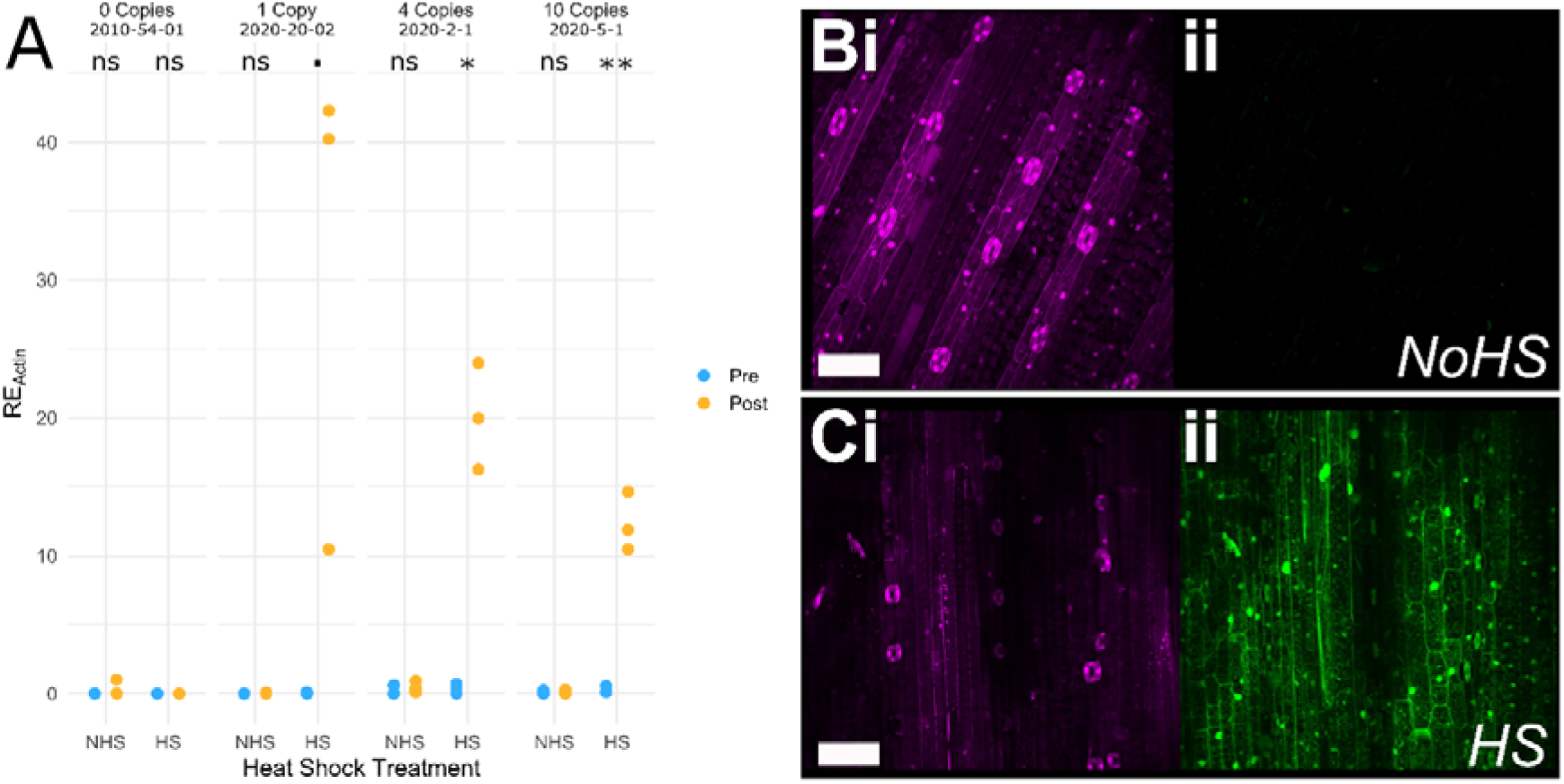
Gene expression is significantly induced following heat shock. (A) Heat shock (HS) at 38 °C is sufficient to induce expression of *NAM-B1* in wheat, while no significant expression is seen in untreated (NHS) plants. (B) No eGFP expression (ii, green) is observed in untreated (No HS) barley seedlings but is visible in all cells following a 30 minute heat shock treatment (C). The reporter gene mCHERRY (magenta) is present in all samples (Bi, Ci). The statistical comparison in panel A was carried out using the Student’s T-test, where N is 3 or 4 for each condition; ns, p > 0.1, ·, p < 0.1, *, p < 0.05, **, p < 0.001.

We then tested whether the heat shock treatment was able to induce expression of *eGFP* in the barley transgenics, using homozygous progeny of the T_0_ line 00899-04-01. A heat treatment of 30 minutes of 38 °C induced high levels of eGFP in the leaf tissue of one-week old seedlings, whereas no background expression of eGFP was observed in the untreated controls (Figure 3B, C). Residual expression of mCHERRY is observed in the leaf tissue following heat shock, though this is substantially reduced compared to the untreated control (Figure 3B, C). This will be due in part to the two copies of the construct present in this line, as excision of only one copy is required to see eGFP signal. The mCHERRY protein is also known to be highly stable (23, 25), and thus some residual protein is likely to remain in the four days between construct excision and tissue imaging.

#### Altered durations of heat shock lead to sectorial induction of the gene of interest

Having demonstrated that application of heat shock is sufficient to induce expression of the gene of interest following excision of the reporter gene, we then investigated whether reducing the duration of heat shock treatment could also cause recombination and gene expression. We visualised the expression of eGFP in 1-week-old barley seedlings following the application of a 38 °C heat shock of varying durations. To assess the effectiveness of the heat shock treatment at penetrating tissue layers, we analysed the eGFP expression in successive leaves from the outer mature leaves, to the innermost leaf primordium (P2) which is completely surrounded by the older leaves.

No expression of eGFP was observed without heat shock, as expected (Figure 4A, B). Five minutes of heat shock was sufficient to induce complete eGFP expression in all outer leaves up to the fourth primordium (P4, Figure 4C). Single cells expressing eGFP were observed in the third primordium (P3) of plants heat-treated for 5 minutes (Figure 4D), while no eGFP was observed in the innermost P2 primordium (Figure 4E). A 15-minute heat treatment, was sufficient to induce eGFP expression in all leaves and tissues, including the second leaf primordium (P2) and the meristem (Figure 4F) in 1-week-old seedlings. This illustrates the tunability of expression for different tissue layers based on heat shock duration.

**Figure 4:**
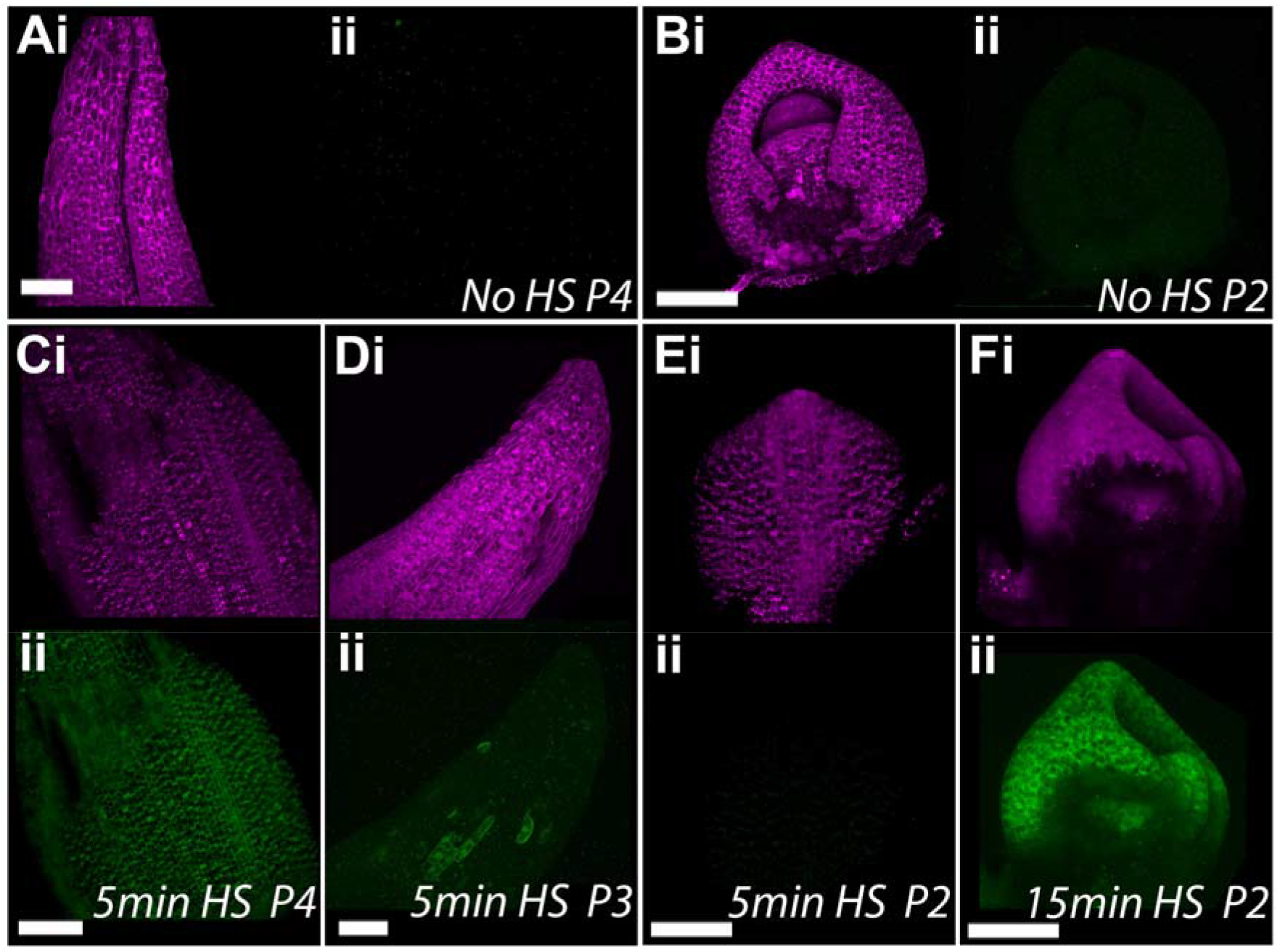
Short-duration heat shock leads to sectorial induction of eGFP. Both the fourth leaf primordia (A) and the second leaf primordia (B) show mCHERRY expression (i) but no eGFP expression (ii) before heat shock. After five minutes of heat shock, all cells in the fourth leaf primordia (C) show eGFP while sectorial expression of eGFP is observed in the third leaf primordia (D). No expression of eGFP is observed in the second leaf primordia after five minutes of heat shock (E). However, after 15 minutes of heat shock, eGFP is expressed in all cells of the meristem and second leaf primordia (F). All plants were imaged three days after treatment. In each pair of images, the reporter gene mCHERRY is shown in panel i (magenta), and eGFP is shown in panel ii (green). Scale bars are 100 μm.

#### Heat shock application at different developmental stages can lead to successful construct activation

After establishing that the applied heat shock was sufficient to induce gene expression in treated seedlings, we next determined the extent to which this was true at later developmental stages. We found that a two-hour application of heat shock at 38 °C at the three-leaf stage in soil-grown barley was sufficient to induce gene expression in the outer leaves (i.e. leaf 3, Figure 5A, B). However, minimal eGFP expression was observed in later developing leaves (i.e. leaf 5, Figure 5C, D) and no eGFP was observed in the inflorescence meristem (Figure 5E, F). This suggests that the heat treatment was not able to fully penetrate to the meristematic cells at the three-leaf stage, resulting in new tissue growth containing only the unexcised construct.

**Figure 5.**
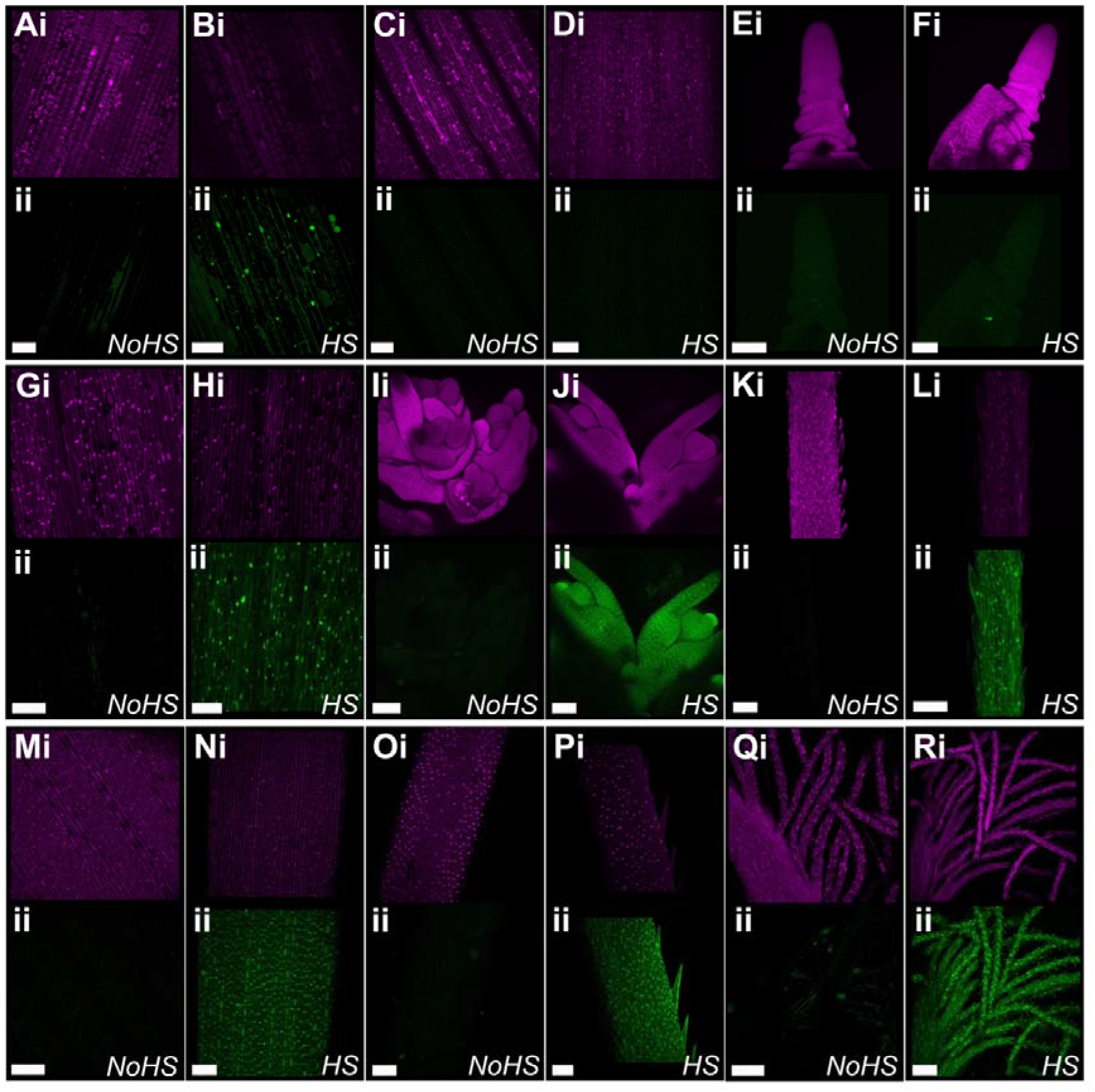
Testing heat shock stages. Heat shock treatments of plants at different developmental stages: 3 leaf stage (A-F), flag leaf emergence (G-L), and ear emergence (M-R). Heat shock was carried out at 38 °C for 2 hours, and each panel shows the mCHERRY expression (i, magenta) and eGFP expression (ii, green). Tissues are imaged in pairs, showing the equivalent tissue without (left) and with (right) heat shock. (A-B) Leaf 3, (C-D) Leaf 5, (E-F) inflorescence meristem. (G-H) flag leaf blade, (I-J) developing florets, (K-L) young awns. (M-N) stem epidermis below the inflorescence, (O-P) mature awn, (Q-R) stigma. Scale bars 100um. At all developmental stages, plants were imaged four days after treatment.

We then investigated whether this result would be recapitulated in the wheat transgenics. Soil-grown plants were treated with a two hour 38 °C heat shock at the three-leaf stage. Similarly to barley, we found that tissue from leaf 3 showed complete excision of the reporter gene (Figure 6A, “3^rd^”), while tissue from leaf 5 showed inconsistent excision between independent replicates (Figure 6A, “5^th^”). Some individuals showed full excision in the leaf 5 tissue (Figure 6A, left panel), indicating that the meristematic cells from which leaf 5 developed had experienced sufficient heat shock for full excision of the reporter gene. Others showed an intermediate state, with the presence of both the full construct, at 2490 bp, and the excised construct, at 151 bp (Figure 6A, middle panel). In these cases, it is likely that the fifth leaf developed from a mixture of excised and unexcised primordia cells, leading to a mosaic in the mature tissue. Finally, some individuals showed no evidence of construct excision, indicating that the fifth leaf primordia cells were not successfully heat shocked (Figure 6A, right panel).

**Figure 6.**
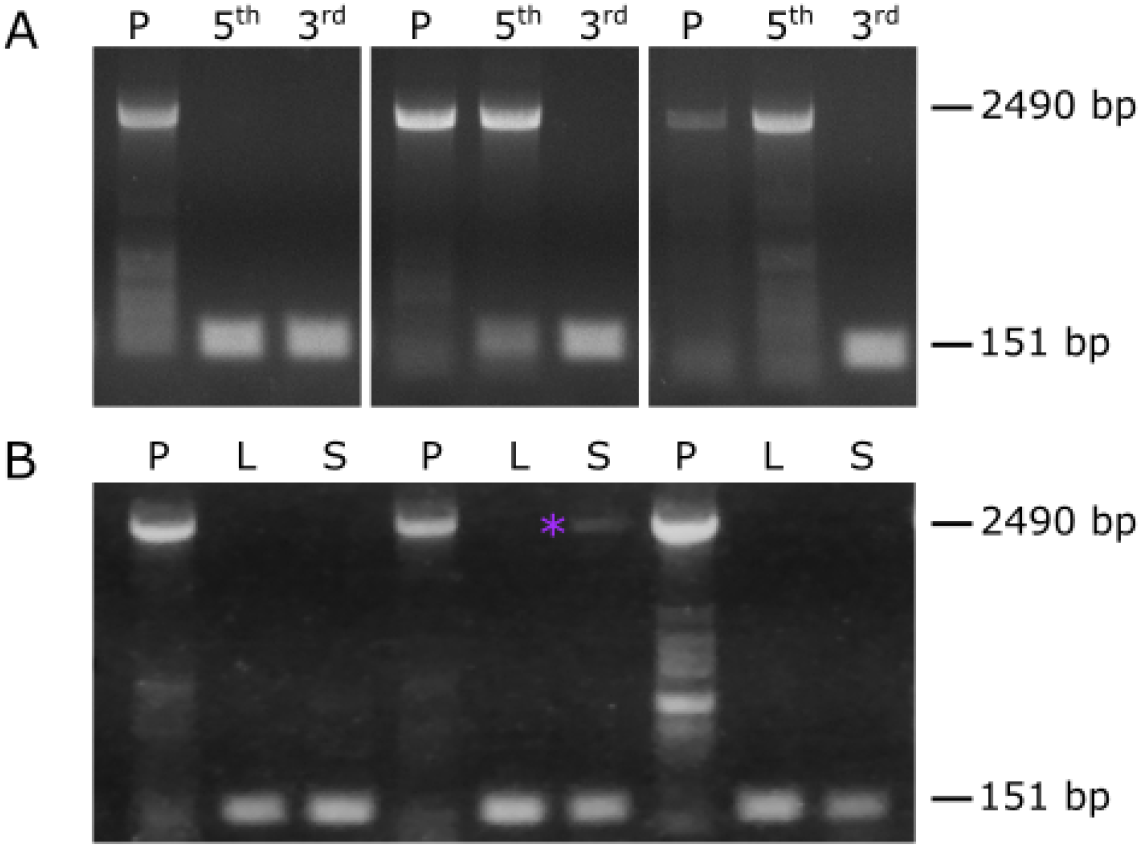
Heat shock treatment does not always lead to complete excision of the reporter gene. At the three-leaf stage (A), two hours of heat shock completely excises the reporter gene from the leaf tissue present at the time of heat shock (third leaf, “3^rd^”; 151 bp). Leaf tissue grown after completion of the heat shock treatment (fifth leaf, “5^th^”) is not always completely excised, as shown by the presence of the reporter gene (2490 bp band). The three panels represent three independent biological replicates. (B) At flag leaf emergence, a two-hour heat shock at 38 °C leads to complete excision in the flag leaf (L) and in the corresponding basal spikelet (S) in most cases, though in some only partial excision is observed in the basal spikelet (purple asterisk). The reporter gene is present in all individuals before heat shock, with no evidence of premature excision (“P”). Gels shown are representative examples from 15 (A) and 12 (B) T_1_ individuals, from lines 2020-20-02, 2020-2-1, and 2020-19-01.

As a two-hour heat shock treatment at 38 °C was not sufficient to consistently induce construct excision in the entirety of the three-leaf stage plant, we decided to investigate whether this treatment would be able to induce construct excision at later developmental stages. At the flag leaf emergence stage, we found that this treatment was sufficient to induce complete construct excision in all aerial tissues in barley (Figure 5G – L). *eGFP* expression was induced in the flag leaf (Figure 5G – H), the developing florets (I – J), and in the young awns (K – L). This was mirrored in the wheat transgenics, where the same heat shock treatment leads to activation of the construct and excision of the reporter gene in both the flag leaf and spikelet tissue (Figure 6B). In some cases, the excision in the basal spikelet was not complete, though the excised construct is also present (Figure 6B; purple asterisk). We also found that the heat shock treatment of the barley plants at spike emergence was also sufficient to induce *eGFP* expression in all aerial organs, including in the stem and the floral organs of the plant (Figure 5M – R).

#### The effect of lower temperatures on construct stability

As treatment at 38 °C can induce expression of the heat-shock driven construct, we then investigated whether temperatures lower than 38 °C could lead to inadvertent (i.e. “leaky”) activation of the construct. We focussed on the barley transgenic system, as the use of fluorescent markers allowed the visualisation of cell-specific changes in expression. In one-week old seedlings, treated with a heat shock of 30 minutes, we only observed high levels of eGFP expression at 38 °C or higher (Figure 7M – T). This was consistent across all three emerged leaves and the meristem. At 32 °C, some individual cells exhibited eGFP expression in the emerged leaves (Figure 7I – K), though no eGFP was observed in the meristem or leaf primordia (Figure 7L). Even fewer cells expressed eGFP at 26 °C, and only in leaves 1 and 2 (Figure 7E – H). No expression of eGFP was observed at 20 °C, in keeping with previous experiments (Figure 7A – D). This was also consistent with previous studies of the *HvHSP17* promoter (10).

**Figure 7:**
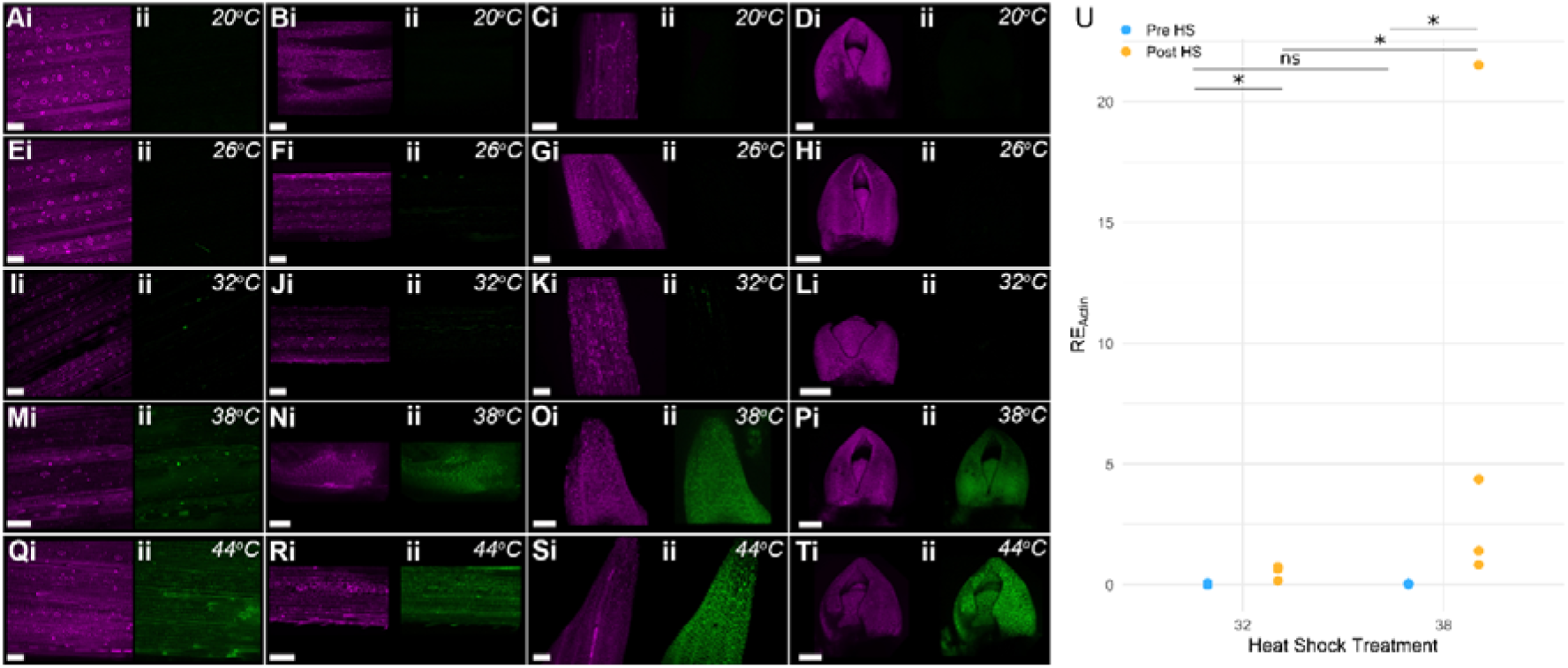
eGFP is strongly induced only at or above 38°C. (A – T) One-week old seedings were subjected to a heat shock of 30 minutes, at one of five different temperatures: 20 °C (A – D), 26 °C (E – H), 32 °C (I – L), 38 °C (M – P), and 44 °C (Q – T). The mCHERRY (i, magenta) and eGFP (ii, green) expression is shown for each tissue sample. Images of leaf 1 (A,E,I,M,Q), leaf 2 (B,F,J,N,R), leaf 3 (C,G,K,O,S), and the meristem with attached leaf primordia (D,H,L,P,T) are shown for each temperature treatment. All plants were imaged three days following treatment. The scale bar is 100 μm. NAM-B1 is induced most strongly at 38 °C, though also to a lesser extent at 32 °C (U). Statistical comparisons were carried out using the Wilcox test due to the non-normality of the data, with N = 4 for all conditions; *, p < 0.05, ns, p > 0.05.

We then tested whether the induction of the heat-shock construct at 32 °C was visible in the wheat transgenic lines, using T_1_ plants descended from the single-copy line 2020-20-02. Ten days after a two-hour heat shock, seedling leaf tissue showed a significant increase in *NAM-B1* expression at 38 °C, as expected (Figure 7U, p < 0.05). Those treated with a two-hour 32 °C heat shock also showed a significant increase in *NAM-B1* expression (Figure 7U, p < 0.05). However, the level of expression at 32 °C was significantly lower than that observed after heat shock at 38 °C (p < 0.05). This mirrors the comparatively low levels of eGFP expression observed at 32 °C in the barley transgenics.

## Discussion

In this paper, we demonstrate a system for inducible and irreversible expression of a gene of interest following application of heat shock. This system is functional at different developmental stages and provides the possibility of complete gene induction in the whole plant, or of sectorial gene induction following a shorter application or lower temperature of heat shock. The successful use of the system will require appropriate monitoring of the construct to ensure that both construct excision has occurred when desired, and that premature excision has not occurred.

### Use in developmental studies

The use of an inducible system such as the one presented here will allow more rigorous and detailed developmental studies in cereal crop species. We have demonstrated that by modulating the length of applied heat shock, it is possible to induce variable levels of construct activation. Single-cell activation can be coupled with imaging of developing organs to effectively study the growth and cell division rates within clonal sectors descended from the activated parent cell (26–28). Similarly, targeted application of heat shock to individual tissues or organs could allow comparison of treated and untreated parts of the plant (29). For example, a single tiller of a developing wheat plant could be covered and exposed to a heat shock in a water bath, while the remainder of the plant is at room temperature and therefore uninduced. Previous work demonstrated that the activation of the *HvHSP17* promoter is highly specific to the treated tissue (10), suggesting that the key limiting factor to this approach would be the ability of the induced protein of interest to travel sym- or apoplastically out of the treated tissue.

This level of specificity compares favourably with that observed in other chemically-inducible systems. Often chemical applications are applied through a foliar spray, in the case of more mature plants, or through growth of seedlings on media containing the chemical inducer (30). Neither of these methods are conducive for tissue-specific induction of the construct, as is possible with the heat shock-inducible system. In some cases, painting of leaves or other tissues with a solution containing the chemical inducer of interest can lead to spatially-specific induction, however this depends on the mobility of the chemical inducer within the cells. Some chemicals, such as 17-beta-oestradiol are relatively immobile, and thus suitable for this approach, while others, such as dexamethasone, are transported symplastically, making them unsuitable for localised studies (30). The application method for chemical inducers also raises difficulties when inducing mature plants. So-called “spray and pray” approaches which rely upon ectopic application of the inducer via a foliar spray are notorious for incomplete and inconsistent induction (31). In contrast, heat shock application at later developmental stages relies upon changing the ambient temperature, which inherently affects the entire exposed surface of the plant.

Inducible systems such as these can also be used to study the effect of overexpressing genes which are known or hypothesized to be embryo lethal. This is particularly important in the case of cereal crops and other species where transformation is still non-trivial and often expensive. By using an inducible system, transformed plants can be successfully recovered at the T_0_ stage without the risk of embryo lethality. After recovering and bulking transformed seeds, the plants can then be heat shocked at the desired developmental stage to induce gene expression. In our case, we initially developed this system in wheat to test the effect of ectopically expressing the wheat senescence regulator *NAM-B1* (24). As a positive regulator of senescence, we hypothesized that this overexpression would be embryo lethal. However, we saw no evidence to suggest that overexpression of *NAM-B1* at seedling stage and pre-anthesis affects plant development or causes any symptoms of premature cell death. Further work is needed to explore why premature expression of *NAM-B1* does not lead to premature senescence or cell death.

This construct can also be adapted for use in more complex systems. For example, the design of the construct could be inverted such that a gene of interest is expressed under the control of the main promoter (in this case *ZmUbi*) and is excised from the construct following heat shock. This setup could be used to allow ectopic expression of a gene only until a specific stage of development, at which point it is removed from the construct and no longer expressed. The system could also be coupled to other transgenic systems, for instance an RNAi (32, 33) or miRNA construct (34) which either begins silencing of the target gene following heat shock, or conversely only silences the targeted gene until heat shock is applied. The potential applications for this system in cereal crops are substantial and are likely to come into wider use as the efficiency of transformation increases and the costs decrease.

### Treatment success depends upon developmental stage

While application of heat-shock to young seedlings was able to rapidly induce complete excision of the reporter gene, within 15 minutes in the case of barley (Figure 4F), such complete excision was more difficult to obtain at the three-leaf stage (Figure 5). Critically, at this stage the plant is sufficiently juvenile that complete excision in the meristem is essential to ensure that the latterly developed tissue is also expressing the gene of interest. A clear distinction between the assays carried out on the seedlings and those carried out on the older plants is that the seedlings were grown on filter paper, while the older plants were grown in soil. As a result, while the meristem of the seedlings was fully exposed to the heat shock, the meristem of the older plants at three-leaf stage would be located underneath the soil. It is possible that the insulation provided by the soil prevents efficient activation of the construct, compared to that seen in aboveground tissues. Notably, we have not characterised the function of this construct in roots, which are likely to be well insulated to external changes in temperature. For those interested in developing this construct for use in roots, we would encourage bespoke optimisation of the heat shock application method. In general, it is essential that the heat shock is applied in such a manner as to ensure the tissue of interest reaches the desired temperature. Where the tissue is insulated from the heat, either by soil, other tissues, or other means, a longer heat shock may be required to induce the desired level of construct activation.

### No construct activation is observed under standard growth conditions

It is essential that researchers can be sure that the inducible promoter will not be prematurely activated at growth temperatures. To ensure that the construct was sufficiently stable at temperatures below 38 °C, we carried out assays at temperatures ranging from 20 °C to 38 °C (Figure 7). While some construct activation was observed at 26 °C and 32 °C, this was significantly less than that observed at 38 °C. Critically, no significant activation of the construct was observed at 20 °C, the standard temperature used under light conditions in our controlled growth environments. This indicates that although there is the possibility of unintended construct activation at temperatures below 38 °C, this should not occur under normal growth conditions. We would therefore strongly encourage those working with these constructs to grow plants in controlled environment conditions where possible, ensuring that the plants remain at or below 20 °C. Where this is not possible (e.g. due to space or facilities constraints), it is essential that the plants are grown alongside ambient temperature monitors, which are regularly monitored for any increases in temperature. In all cases, plants should be tested at critical experimental junctures for any unintentional activation of the construct, most cost-effectively through PCR-based assays of construct excision (e.g. in Figure 2). Experiments should be designed with redundancy in the plant materials, to allow for the loss of individual plants if undesirable construct expression is detected. Finally, and perhaps most critically given the irreversible nature of the excision, seed stocks of lines containing the inactivated construct must be stored with the utmost care to ensure that they remain inactivated. Ideally cold storage would be used, but at a minimum, seeds should be stored away from any sources of ambient heat i.e. sunny windows or radiators, which could heat the seeds above 20 °C.

## Conclusions

Here we have described a heat-shock inducible system which drives constitutive gene expression in barley and wheat. This system, using standard Golden Gate constituent parts, can be adapted for use in developmental studies of genes which may otherwise be recalcitrant to transformation. It can also be used to study cell-level development and could be adapted for use in many other contexts. We have shown that the system is versatile, stable and functional in multiple wheat and barley tissues and at various developmental stages. As with all systems of this kind, care must be taken to ensure that construct activation only occurs when desired, and that is has occurred to the expected extent. As more cereal research moves towards gene functional characterisation at the molecular and cellular level (1), transgenic approaches such as this will become increasingly relevant.

## Materials and Methods

### Construct design

Golden-gate assembly (35, 36) was used to develop all constructs, utilising parts from TSL SynBio (synbio.tsl.ac.uk) and ENSA (ensa.ac.uk) and following the standard Type IIS cloning syntax, where possible (37). All constructs used in the course of this study are listed in Supplementary Table 2.

Novel parts were created using DNA synthesis using domestication to remove any endogenous *Bsa*I, *Esp*3I, and *Bpi*I sites from published sequences using neutral base pair changes. The Cre recombinase gene sequence (Genbank GeneID: 2777477) and *loxP* sites (5⍰ – GACCTAATAACTTCGTATAGCATACATTATACGAAGTTATATTAAGGGTTG- 3⍰) were taken from the genome sequence of Enterobacteria Phase P1, domesticated where necessary, and synthesized (Invitrogen Gene Systems).The functional *NAM-B1* coding sequence (from *Triticum turgidum* ssp. *dicoccoides*) was obtained from NCBI (GenBank DQ869673.1) and domesticated to remove the single *Bbs*I site present in exon 3 (Supplementary File 1). Introns were approximated based on the gene sequence of the functional homoeolog *NAM-A1* in the Chinese Spring reference sequence (TGACv1) (38). The domesticated gene, with introns, was synthesized by Genewiz.

#### Level 0 construct synthesis

To prevent expression of the Cre recombinase during cloning in *E. coli*, we introduced the *Arabidopsis thaliana* U5 small nuclear ribonucleoprotein component intron (from pICSL80006, TSL SynBio) at 254 bp in the coding sequence, based on published mammalian studies (39). The Cre recombinase gene was amplified in two sections, using primer sets designed using the general format: NGAAGACNN + *Bpi*I-specific 4 bp overhang + 18-30bp of the target sequence ((37); Supplementary Table 3), which introduced flanking *Bpi*I sites and allowed ligation into the pICH41308 backbone. Simultaneously, the U5 intron sequence was amplified (Supplementary Table 2) to allow ligation between the two Cre recombinase sequence fragments. 2μl of each PCR fragment was included in the ligation reaction, following the protocol described in (35). The resulting sequence is provided in Supplementary File 1.

The domesticated *NAM-B1* sequence, and the sequence of the ZmUbi promoter, along with its 5’UTR (from pICSL12009) were cloned directly into the universal acceptor pUAP1 (37) to generate level 0 constructs with the desired 4 base pair overhangs, hereafter pNAM-B1 and pZmUbi, respectively (see primers used in Supplementary Table 3). The PCR-amplified sequences were ligated into pUAP1 using the *Bpi*I enzyme, following the protocol from (35).

#### Level 0.5 construct synthesis

To introduce flanking *loxP* sites around the reporter gene and corresponding terminator, an additional step was required alongside the standard Golden Gate cloning protocol. The short *Bsa*I protocol was carried out as described by TSL Synbio, using the *loxP* level 0.5 construct developed by ENSA (EC10161) (Figure 8A). For the wheat construct, the GUS coding sequence from TSL Synbio (pICH75111) and the nosT terminator sequence (pICH41421) were cloned into the *loxP* construct (Supplementary Figure 2A). For the barley construct, an ER targeting sequence (EC71090), the mCHERRY coding sequence (EC71091), the HDEL sequence (EC71020), and the 35S terminator (EC41414) were cloned into the *loxP* construct (Supplementary Figure 3A). This created a “Level 0.5” vector, hereafter referred to as p*loxP*-GUS or p*loxP*-mCHERRY, with *loxP* sites flanking the coding region and terminator. Unlike standard Level 0 constructs, the Level 0.5 constructs contain flanking *Esp*3I sites, rather than *Bsa*I sites. The Level 0.5 construct combines positions B1 and B2 of the standard Type IIS cloning syntax, allowing its insertion upstream of standard coding regions, and downstream of standard promoter regions (37).

**Figure 8:**
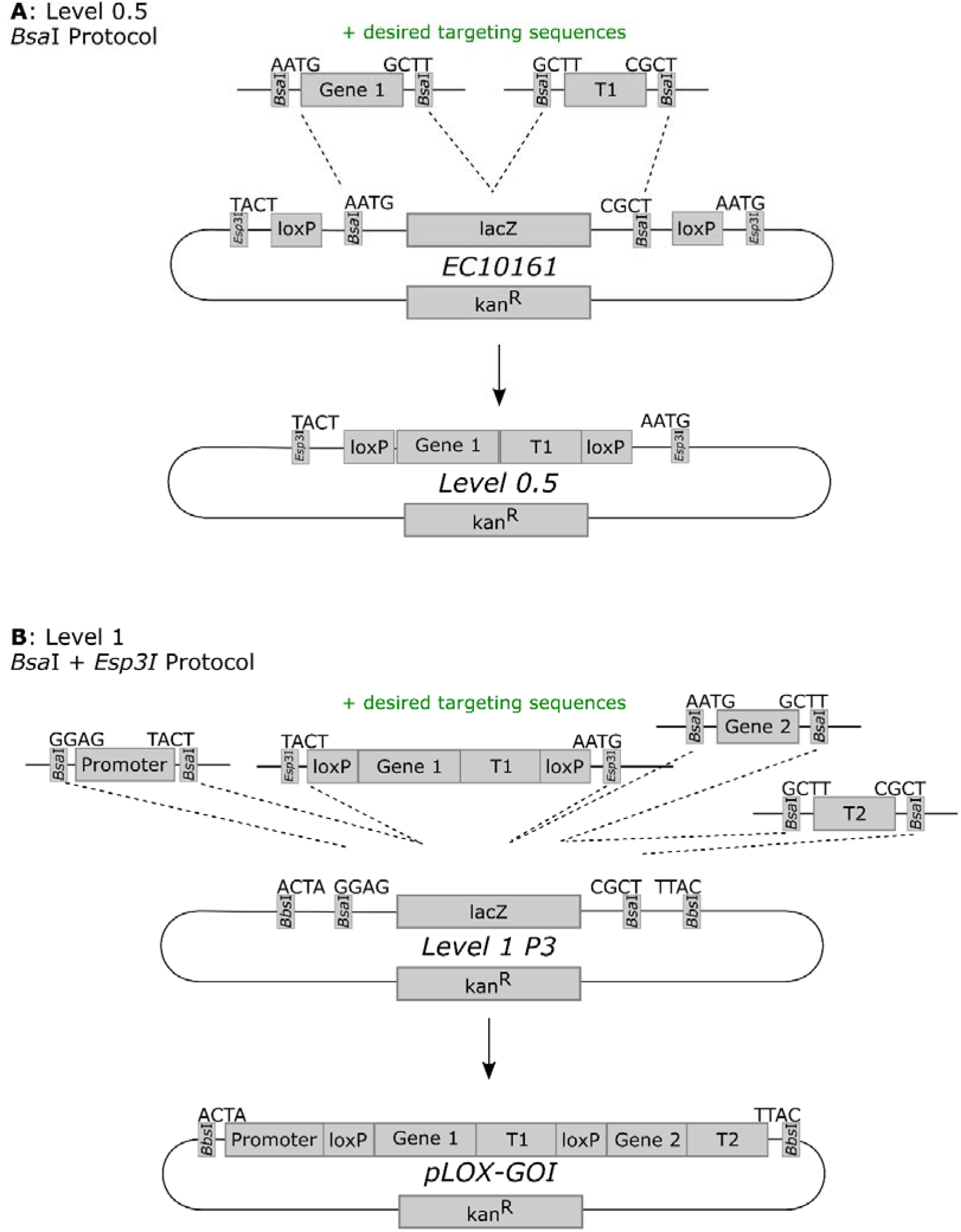
Assembly of the Level 0.5 and Level 1 vectors. (A) The reporter gene (Gene 1) and its associated terminator (“T1”) were cloned into the Level 0 vector EC10161, using the Type IIS restriction enzyme *Bsa*I. The resulting Level 0.5 vector contains *loxP* sites flanking the inserted reporter gene and terminator. (B) The desired promoter sequence, *loxP*-flanked reporter gene, gene of interest (Gene 2), and corresponding terminator (T2) are cloned into a Level 1, position 3 vector (e.g. pAGM8031) in a reaction involving both *Bsa*I, for the canonical Level 0 and Level 1 parts, and *Esp*3I, for the *loxP*-containing Level 0.5 vector (see A). In both cases, additional components can be included during the cloning step, such as targeting sequences.

#### Level 1 construct synthesis

For each system, a level 1 construct was synthesized that combined a constitutive promoter, the *loxP*-flanked reporter gene, and the gene of interest (Figure 8B). For the wheat construct, the promoter construct (pZmUbi), the *loxP* construct (p*loxP*-GUS), the gene of interest (pNAM-B1), and the nosT terminator (pICH41421) were then cloned into the Level 1 backbone, position 3 (pICH47751) (Supplementary Figure 2B). For the barley construct, the promoter construct (EC71139), the *loxP* construct (p*loxP*-mCHERRY), the ER targeting sequence (EC71090), the gene of interest (EC71088), the HDEL sequence (EC71020), and the actin terminator (EC44300) were then cloned into a Level 1 backbone, position 3 (EC47822) (Supplementary Figure 3B). In both cases, this step used a modified version of the *Bsa*I protocol from (35) which contains 10 U *Bsa*I and 10 U *Esp*3I, instead of 20 U of *Bsa*I, to allow insertion of the *loxP* construct (Figure 1B).

#### Level 2/M construct synthesis

The level 1 constructs were cloned into the appropriate final backbone (Level 2/M), integrating the *loxP* construct with the Cre recombinase construct, and the Hygromycin selection cassette, flanked by right and left border repeats for *Agrobacterium*-mediated transformation. For the wheat construct, the NAM-B1-*loxP* level 1 construct was cloned into the level M backbone (pAGM8031), alongside a selection cassette (OsAct∷Hygromycin, pICH47802), the HvHsp17∷Cre construct (EC71173), and the appropriate end linker (pICH50892; Supplementary Figure 2C). For the barley construct, the level 1 eGFP construct, the HvHSP17∷Cre construct (EC71173), and the appropriate end linker (EC41766) were cloned into the level 2 plasmid EC15027, which contains an appropriate Hygromycin expression system (Supplementary Figure 3C). Cloning was carried out using the *Bbs*I protocol from (35). These constructs will be referred to as HS_NAM-B1 and HS_GFP, respectively. The sequences of all constructs at all stages were verified using Sanger sequencing (Eurofins). The specific cloning steps for the wheat and barley constructs, respectively, are illustrated in Supplementary Figures 2 and 3.

### Transformation and copy number determination

The wheat construct was transformed into *Triticum aestivum* cv. Fielder following previously established protocols as detailed in (2). The barley construct was transformed into *Hordeum vulgare* cv. Golden Promise (40). Transformation was carried out by the BRACT platform at the John Innes Centre. Positive transformants were identified at the T_0_ generation by iDNA Genetics (Norwich, UK) using a Taqman probe against the Hygromycin resistance cassette, as described in (2). This process also assigned copy numbers of the construct to each independent event. For the barley construct, copy number analysis was also repeated at the T_1_ generation to identify stable homozygous transformants for use in microscopy.

### Wheat growth and phenotyping

#### Seedling heat shock assays

Seeds were germinated at 4 °C on damp filter paper for 48 hours in the dark before they were moved to room temperature and were grown under ambient light conditions. The seedlings were grown for five to seven days following germination before the heat shock treatment. Half of the seedlings were treated to a two-hour heat shock at 38 °C in a temperature-controlled Ecotron incubator (Infors HT). The remaining seedlings were left at ambient room temperature during this period.

#### Controlled environment experiments

Seeds were germinated at 4 °C in the dark for 48 hours on damp filter paper, and then allowed to grow for a further 48 hours under ambient light conditions. The germinated seedlings were then sown into P96 trays containing 85% fine peat with 15% horticultural grit. They were grown until the 2-3 leaf stage before transplanting into 1 L pots. The plants experienced cycles of 16 hours of light at 20 °C and 8 hours of dark at 15 °C.

Where required, a heat shock treatment was applied to plants grown in soil. Depending on the developmental stage of the plant, either a P96 tray (3-leaf stage or earlier) or up to nine 1 L pots (flag leaf emergence or later) were taken to a controlled environment chamber (Sanyo MLR-351H) and incubated at 38 °C for two hours, at 85% relative humidity and light level of 3 lumens, before being returned to the original environment. A second set of non-heat shock controls were left in the original environment for the period of the heat shock.

### Barley growth and phenotyping

#### Standard Growth conditions

T_0_ plants were grown in 1 L pots in John Innes cereal mix in controlled environment rooms (Gallen Kamp) at 75% humidity, with a day/night cycle of 16 hours light at 15 °C, 8 hours dark at 12 °C. T_1_-T_3_ generations were grown in 1 L pots of cereal mix in temperature-controlled greenhouse conditions, at 20 °C during the day and 15 °C during the night, and allowed to self to bulk seed.

#### Heat shock experiments

##### 1-week-old seedlings

Seeds were germinated on filter paper in plates at room temperature and grown for 1 week before treatment. For heat shock treatment, the seedlings were placed into pre-warmed plastic wallets which were sealed before being immersed in a water bath set to the desired temperature for the desired length of time. Following treatment, the seedlings were removed from the wallets and plated on filter paper to grow for three days before dissection and imaging.

##### Soil-grown plants

Seeds were germinated at 4 °C in the dark for 48 hours on damp filter paper, and then allowed to grow for a further 48 hours at room temperature in the dark. The seedlings were then sown in P96 trays containing M3 mix with insecticide, and grown under controlled conditions with 16 hours light, at 20° C and 8 hours of dark, at 15° C. Plants treated at the three-leaf stage were heat shocked in an incubator for 2 hours at 38 °C while in the P96 trays. Those treated at the flag leaf emergence and ear emergence stages were transplanted to 1 L pots before receiving heat shock treatment in the incubator, again at 38 °C for two hours. In all cases, plants were returned to the controlled environment conditions for four days before dissection and imaging.

### Construct Excision PCR

To determine whether the application of heat shock was able to induce excision of the *loxP*-flanked region by the Cre recombinase, we carried out PCR on genomic DNA (gDNA). gDNA was extracted using the standard DNA extraction protocol, as described in (41). Primers which flanked the *loxP* sequences were designed (42) which would amplify different sized fragments depending on the reporter gene being excised or not (Figure 2A, Supplementary Table 3**Error! Reference source not found.**). The PCR was carried out under the following conditions: 5 min at 95 °C, 10 cycles of 30 seconds at 95 °C, 1 minute starting at 63 °C and decreasing by 1 °C per cycle, and 3 minutes at 68 °C, a further 35 cycles of 30 seconds at 95 °C, 1 minute at 53 °C, and 3 minutes at 68 °C, followed by a final extension at 68 °C.

### qPCR

#### RNA extractions

Leaf tissue was sampled from individual plants and snap frozen in liquid N_2_. The stage at which the tissue was sampled depended on the experiment in question. Initial validation of construct expression was carried out on seedling leaf samples, taken between five- and seven-days following germination from the first true leaf of the plant (i.e. second leaf stage). Tissue samples from plants grown on soil in controlled environment conditions were taken at the third leaf stage. In all cases, approximately 3 cm of tissue were sampled from the tip of the leaf. The snap-frozen tissue was then ground to a fine powder and RNAs were extracted using TRIzol^®^ Reagent (ThermoFisher). cDNA synthesis was carried out using the Invitrogen M-MLV reverse transcriptase, following treatment with RQ1 RNase-Free DNase (Promega).

#### RT-PCR

Initial validation of Cre recombinase expression was carried out using RT-PCR on cDNA samples from lines carrying the heat-shock construct (HS_NAM-B1). Touchdown PCR was carried out on the relevant samples using NEB *Taq* polymerase using the primer pair SH002/SH053 (Supplementary Table 3) under the following conditions: 5 min at 95° C, 10 cycles of 30 seconds at 95°C, 1 minute starting at 62° C and decreasing by 1° per cycle, and 1 minute at 68 °C, a further 25 cycles of 30 seconds at 95 °C, 1 minute at 52° C, and 1 minute at 68° C, followed by a final extension at 68° C.

#### qRT-PCR

The expression of *NAM-B1* in the wheat transgenic lines was validated using quantitative RT-PCR. Primers specific to the domesticated version of *NAM-B1* present in the construct were designed and used (SH049/SH051) alongside previously published primers for the internal control gene *TaActin* ((24); Supplementary Table 3). Efficiencies for the primers were calculated using pooled cDNA from samples which were known to be expressing the transgenic construct (Supplementary Table 3). qRT-PCR reactions were performed using the LightCycler^®^ 480 SYBR Green I Master Mix with a LightCycler 480 instrument (Roche Applied Science, UK) under the following conditions: 5 min at 95 °C; 45 cycles of 10 s at 95 °C, 15 s at 60 °C, 30 s at 72 °C; followed by a dissociation curve from 60 °C to 95 °C to determine primer specificity. In all cases, three technical replicates were carried out per sample and the expression of *NAM-B1* was recorded relative to *TaActin*.

### Microscopy

The spectral scan was carried out using an SP5 II Leica confocal laser microscope, with a x20 water immersion lens. During imaging the gain was increased until signal could be seen in each of the fluorescent marker channels (CyPET: 449-508nm, eGFP: 518-565nm, mCHERRY: 586-644nm). In all wild-type tissues this was very high, with gains of over 150 required to visualise the autofluorescence.

Imaging of the heat shocked plants was carried out on a Leica SP8 confocal microscope with a x20 water immersion objective. Laser power was set to 1 % and HyD detectors were used for both the mCHERRY and eGFP, with gain <75 and pinhole set to 1AU. Z-stack images were collected with line sequential scans to minimise crosstalk between the mCHERRY and eGFP emissions. Images were analysed using FIJI (43) and figures were generated using Adobe Photoshop.

### Statistical Analysis and Data Visualisation

All statistical tests were carried out in R (v 3.6.3). Where data followed the normal distribution, the Student’s T-test was used to compare samples, and where this assumption was not met, the non-parametric Wilcoxon Rank-Sum test was applied. The specific test used in each case is noted within the results and the figure captions. Sample size is indicated in figure captions. All graphs were created in R, using ggplot2 (44), and statistical comparisons were added using Inkscape.

## Declarations

### Additional Files

Supplementary File 1 (.docx) contains gene sequences of *NAM-B1* and *Cre* which were domesticated for use in cloning. Supplementary File 2 (.docx) contains all Supplementary Figures and Tables referred to in the text.

### Availability of Data and Materials

All constructs generated in this study will be made available on Addgene (https://www.addgene.org/Cristobal_Uauy/). Deposition and validation of constructs has been delayed due to the outbreak of COVID-19, however these will be made available as soon as possible. Constructs used in this study which were developed elsewhere are listed in Supplementary Table 1 and can be accessed through contacting the appropriate institution. Note that ENSA has no facility for directly distributing constructs, however ENSA will put interested researchers in contact with those who can distribute the constructs. Germplasm generated in this study is also available from corresponding authors.

### Author’s Contributions

AR, CU, SAH, and PB conceived the study. AR, SAH, CR, and SF carried out the construct design and cloning. AR characterised the barley construct. SAH and AB characterised the wheat construct. SAH drafted the manuscript, and SAH, AR, and CU edited the manuscript. All authors read and approved the final manuscript.

## Acknowledgements

We gratefully acknowledge the support from BRACT at the JIC for carrying out transformation and selection of the transgenic barley and wheat plants, and the support from the Horticultural Services at the John Innes Centre while growing the transgenic lines. We also acknowledge support provided by the University of Edinburgh Plant Growth facilities, in particular Dr. Sophie Haupt and Patricia Watson. We also acknowledge the resources provided from TSL SynBio and ENSA, particularly for providing easy access to their constructs and support in the design and cloning stages. We also acknowledge Professor Enrico Coen at the John Innes Centre for providing initial financial support when developing the system.

## References

1. Adamski NM, Borrill P, Brinton J, Harrington SA, Marchal C, Bentley AR, et al. A roadmap for gene functional characterisation in crops with large genomes: Lessons from polyploid wheat. eLife. 2020; 9:e55646.

2. Hayta S, Smedley MA, Demir SU, Blundell R, Hinchliffe A, Atkinson N, et al. An efficient and reproducible Agrobacterium-mediated transformation method for hexaploid wheat (*Triticum aestivum* L.). Plant Methods. 2019; 15(1):121.

3. Caddick MX, Greenland AJ, Jepson I, Krause KP, Qu N, Riddell KV, et al. An ethanol inducible gene switch for plants used to manipulate carbon metabolism. Nat Biotechnol. 1998; 16(2):177–80.

4. Bruce W, Folkerts O, Garnaat C, Crasta O, Roth B, Bowen B. Expression profiling of the maize flavonoid pathway genes controlled by estradiol-inducible transcription factors CRC and P. Plant Cell. 2000; 12(1):65–80.

5. Zuo J, Niu QW, Chua NH. Technical advance: An estrogen receptor-based transactivator XVE mediates highly inducible gene expression in transgenic plants. Plant J. 2000; 24(2):265–73.

6. Roslan HA, Salter MG, Wood CD, White MR, Croft KP, Robson F, et al. Characterization of the ethanol-inducible alc gene-expression system in *Arabidopsis thaliana*. Plant J. 2001; 28(2):225–35.

7. Kirch T, Simon R, Grunewald M, Werr W. The DORNROSCHEN/ENHANCER OF SHOOT REGENERATION1 gene of Arabidopsis acts in the control of meristem ccll fate and lateral organ development. Plant Cell. 2003; 15(3):694–705.

8. Jasinski S, Piazza P, Craft J, Hay A, Woolley L, Rieu I, et al. KNOX action in Arabidopsis is mediated by coordinate regulation of cytokinin and gibberellin activities. Curr Biol. 2005;15(17):1560–5.

9. Shinmyo A, Shoji T, Bando E, Nagaya S, Nakai Y, Kato K, et al. Metabolic engineering of cultured tobacco cells. Biotechnol Bioeng. 1998; 58(2-3):329–32.

10. Freeman J, Sparks CA, West J, Shewry PR, Jones HD. Temporal and spatial control of transgene expression using a heat-inducible promoter in transgenic wheat. Plant Biotechnol J. 2011; 9(7):788–96.

11. Kovalchuk N, Jia W, Eini O, Morran S, Pyvovarenko T, Fletcher S, et al. Optimization of *TaDREB3* gene expression in transgenic barley using cold-inducible promoters. Plant Biotechnol J. 2013; 11(6):659–70.

12. Xue GP, Way HM, Richardson T, Drenth J, Joyce PA, McIntyre CL. Overexpression of *TaNAC69* leads to enhanced transcript levels of stress up-regulated genes and dehydration tolerance in bread wheat. Mol Plant. 2011; 4(4):697–712.

13. Sauer B. Inducible gene targeting in mice using the Cre/*lox* system. Methods. 1998; 14(4):381–92.

14. Dale EC, Ow DW. Intra- and intermolecular site-specific recombination in plant cells mediated by bacteriophage P1 recombinase. Gene. 1990; 91(1):79–85.

15. Odell J, Caimi P, Sauer B, Russell S. Site-directed recombination in the genome of transgenic tobacco. Mol Gen Genet. 1990; 223(3):369–78.

16. Srivastava V, Anderson OD, Ow DW. Single-copy transgenic wheat generated through the resolution of complex integration patterns. PNAS. 1999; 96(20):11117.

17. Joubès J, De Schutter K, Verkest A, Inzé D, De Veylder L. Conditional, recombinase-mediated expression of genes in plant cell cultures. Plant J. 2004; 37(6):889–96.

18. Sieburth LE, Drews GN, Meyerowitz EM. Non-autonomy of AGAMOUS function in flower development: use of a Cre/loxP method for mosaic analysis in Arabidopsis. Development. 1998; 125(21):4303–12.

19. Gallois J-L, Woodward C, Reddy GV, Sablowski R. Combined SHOOT MERISTEMLESS and WUSCHEL trigger ectopic organogenesis in *Arabidopsis*. Development. 2002; 129(13):3207.

20. Bencivenga S, Serrano-Mislata A, Bush M, Fox S, Sablowski R. Control of Oriented Tissue Growth through Repression of Organ Boundary Genes Promotes Stem Morphogenesis. Dev Cell. 2016; 39(2):198–208.

21. Nguyen AW, Daugherty PS. Evolutionary optimization of fluorescent proteins for intracellular FRET. Nat Biotechnol. 2005; 23(3):355–60.

22. Zhang G, Gurtu V, Kain SR. An enhanced green fluorescent protein allows sensitive detection of gene transfer in mammalian cells. Biochem Biophys Res Commun. 1996; 227(3):707–11.

23. Shaner NC, Campbell RE, Steinbach PA, Giepmans BN, Palmer AE, Tsien RY. Improved monomeric red, orange and yellow fluorescent proteins derived from *Discosoma* sp. red fluorescent protein. Nat Biotechnol. 2004; 22(12):1567–72.

24. Uauy C, Distelfeld A, Fahima T, Blechl A, Dubcovsky J. A NAC Gene regulating senescence improves grain protein, zinc, and iron content in wheat. Science. 2006; 314(5803):1298–301.

25. Hebisch E, Knebel J, Landsberg J, Frey E, Leisner M. High variation of fluorescence protein maturation times in closely related Escherichia coli strains. PLoS One. 2013; 8(10):e75991-e.

26. Kuchen EE, Fox S, de Reuille PB, Kennaway R, Bensmihen S, Avondo J, et al. Generation of leaf shape through early patterns of growth and tissue polarity. Science. 2012; 335(6072):1092–6.

27. Sauret-Güeto S, Schiessl K, Bangham A, Sablowski R, Coen E. JAGGED controls Arabidopsis petal growth and shape by interacting with a divergent polarity field. PLoS Biol. 2013; 11(4):e1001550-e.

28. Whitewoods CD, Gonçalves B, Cheng J, Cui M, Kennaway R, Lee K, et al. Evolution of carnivorous traps from planar leaves through simple shifts in gene expression. Science. 2020; 367(6473):91.

29. Serrano-Mislata A, Bencivenga S, Bush M, Schiessl K, Boden S, Sablowski R. DELLA genes restrict inflorescence meristem function independently of plant height. Nat Plants. 2017; 3(9):749–54.

30. Moore I, Samalova M, Kurup S. Transactivated and chemically inducible gene expression in plants. Plant J. 2006; 45(4):651–83.

31. Kuhlemeier C, Green P. Studying plant development: an alternative to “spray and pray”. Genes Dev. 1987; 1(1):3–5.

32. Gupta S, Schoer RA, Egan JE, Hannon GJ, Mittal V. Inducible, reversible, and stable RNA interference in mammalian cells. PNAS. 2004; 101(7):1927.

33. Tiscornia G, Tergaonkar V, Galimi F, Verma IM. CRE recombinase-inducible RNA interference mediated by lentiviral vectors. PNAS. 2004; 101(19):7347.

34. Garwick-Coppens SE, Herman A, Harper SQ. Construction of permanently inducible miRNA-based expression vectors using site-specific recombinases. BMC Biotechnol. 2011; 11:107–.

35. Weber E, Engler C, Gruetzner R, Werner S, Marillonnet S. A modular cloning system for standardized assembly of multigene constructs. PLoS One. 2011; 6(2):e16765.

36. Werner S, Engler C, Weber E, Gruetzner R, Marillonnet S. Fast track assembly of multigene constructs using Golden Gate cloning and the MoClo system. Bioengineered. 2012; 3(1):38–43.

37. Patron NJ, Orzaez D, Marillonnet S, Warzecha H, Matthewman C, Youles M, et al. Standards for plant synthetic biology: a common syntax for exchange of DNA parts. New Phytol. 2015; 208(1):13–9.

38. Clavijo BJ, Venturini L, Schudoma C, Accinelli GG, Kaithakottil G, Wright J, et al. An improved assembly and annotation of the allohexaploid wheat genome identifies complete families of agronomic genes and provides genomic evidence for chromosomal translocations. Genome Res. 2017; 27(5):885–96.

39. Kaczmarczyk SJ, Green JE. A single vector containing modified cre recombinase and LOX recombination sequences for inducible tissue-specific amplification of gene expression. Nucleic Acids Res. 2001; 29(12):e56–e.

40. Harwood WA. A protocol for high-throughput Agrobacterium-mediated barley transformation. Methods Mol Biol. 2014; 1099:251–60.

41. Pallotta MA, Warner P, Fox RL, Kuchel H, Jefferies SJ, Langridge P, editors. Marker assisted wheat breeding in the southern region of Australia. 10th International Wheat Genetics Symposium; 2003; Paestum, Italy: Instituto Sperimentale per la Cerealicolutra.

42. Borrill PGM. The NAM-B1 transcription factor and the control of grain composition in wheat.: University of East Anglia; 2013.

43. Schindelin J, Arganda-Carreras I, Frise E, Kaynig V, Longair M, Pietzsch T, et al. Fiji: an open-source platform for biological-image analysis. Nat Methods. 2012; 9(7):676–82.

44. Wickham H. ggplot2: Elegant Graphics for Data Analysis. New York: Springer-Verlag; 2016.

