## Supplementary File 1 for "A Heat-Shock Inducible System for Flexible Gene Expression in Cereals"

Supplementary File 1: Gene Sequences

Domesticated *NAM-B1* sequence

The TtNAM-B1 gene sequence was domesticated to remove any *Bbs*I or *Bsa*I sites. One *Bbs*I site was identified (highlighted in yellow in the third exon), and was domesticated from “GTCTTC” to “ATCGTC”, removing the enzyme cut site but retaining the correct amino acid sequence. Exons are coloured, with the splice sites highlighted in red. The exonic sequence is based on the sequence of *NAM-B1* from *T. turgidum* ssp. *dicoccoides*, while the intronic sequence is taken from the non-functional copy of *NAM-B1* present in Chinese Spring.

>full-length domesticated *NAM-B1*

**ATG**GGCAGCTCCGACTCATCTTCCGGCTCGGCGCAAAAAGCAACGCGGTATCACCATCAGCATCAGCCGCCGCCTCCGCAGCGGGGCTCGGCGCCGGAGCTCCCGCCGGGCTTCCGGTTCCACCCGACGGACGAGGAGCTGGTGGTGCACTACCTCAAGAAGAAGGCCGACAAGGCGCCGCTCCCCGTCAACATCATCGCCGAGGTGGATCTCTACAAGTTCGACCCATGGGAGCTCCCCGGTATGTTATGTCTATCTCGTCGGCCGGCCGTGCTTACTTTATCAAGCGCCGCAAATTTTAGGTGCAATTAAATAATCGAATAATCCATCCATCTCATGCTTATACTCCTGTGCACAAGTAGTATTTTTATATTCTTCCAGTACACATGTGTGTAGATGGTTTATGTATGTGATCCTGTCGTGCTTGTTCATGCGCTCGGGATCCGGATCCATCAGAGAAGGCGACCATCGGGGAGCAGGAGTGGTACTTCTTCAGCCCGCGCGACCGCAAGTACCCCAACGGCGCGCGGCCGAACCGGGCGGCGACGTCGGGGTACTGGAAGGCCACCGGCACGGACAAGCCTATCCTGGCCTCGGGGACGGGGTGCGGCCTGGTCCGGGAGAAGCTCGGCGTCAAGAAGGCGCTCGTGTTCTACCGCGGGAAGCCGCCCAAGGGCCTCAAAACCAACTGGATCATGCACGAGTACCGCCTCACCGACGCATCTGGCTCCACCACCGCCACCAACCGACCGCCGCCGGTGACCGGCGGGAGCAGGGCTGCTGCCTCTCTCAGGGTACGTACACGTGTCGATCGCACGGTCTAGCAGTATTTAATTGCTCTCCAGCTTAATTAGGGTATTGTTGATGGTTGATGAAGTTGGACGACTGGGTGCTGTGCCGCATCTACAAGAAGATCAACAAGGCCGCGGCCGGCGATCAGCAGAGGAACACGGAGTGCGAGGACTCCGTGGAGGACGCGGTCACCGCGTACCCGCTCTATGCCACGGCGGGCATGACCGGTGCAGGTGCGCATGGCAGCAACTACGCTTCACCTTCACTGCTCCATCATCAGGACAGCCATTTCCTGGACGGCCTGTTCACAGCAGACGACGCCGGCCTCTCGGCGGGCGCCACCTCGCTGAGCCACCTAGCAGCGGCGGCGAGGGCAAGCCCGGCTCCGACCAAACAGTTTCTCGCCCCGTCATCGTCGACCCCGTTCAACTGGCTCGATGCGTCACCAGTCGGCATCCTCCCACAGGCAAGGAATTTTCCTGGGTTTAACAGGAGCAGAAACGTCGGCAATATGTCGCTGTCATCGACGGCCGACATGGCTGGCGCAGTGGACAACGGTGGAGGCAATGCGGTGAACGCCATGTCTACCTATCTTCCCGTGCAAGACGGGACGTACCATCAGCAGCATGTCATCCTCGGCGCTCCGCTGGTGCCAGAAGCCGCCGCCGCCACCTCTGGATTCCAGCATCCCGTTCAAATATCCGGCGTGAACTGGAATCCC**TGA**

Domesticated Cre-U5-Cre recombinase sequence

The Cre recombinase sequence was domesticated to remove any *Bbs*I or *Bsa*I sites. One BbsI restriction site was identified, and both domesticated from “CTTCTG” to “CTCCTG” via a silent T/C mutation. The U5 intron sequence (purple) was obtained from pICSL80006 (TSL SynBio), with exons coloured and splice sites highlighted in red.

>NC_005856.1:436-1467 Enterobacteria phage P1, complete genome

atgtccaatttactgaccgtacaccaaaatttgcctgcattaccggtcgatgcaacgagtgatgaggttcgcaagaacctgatggacatgttcagggatcgccaggcgttttctgagcatacctggaaaatgctcctgtccgtttgccggtcgtgggcggcatggtgcaagttgaataaccggaaatggtttcccgcagaacctgaagatgttcgcgattatcttctatatcttcaggtaagtttctgcttctacctttgatatatatataataattatcattaattagtagtaatataatatttcaaatatttttttcaaaataaaagaatgtggtatatagcaattgcttttctgtagtttataagtgtgtatattttaatttataacttttctaatatatgaccaaaatttgttgatgtgcaggcgcgcggtctggcagtaaaaactatccagcaacatttgggccagctaaacatgcttcatcgtcggtccgggctgccacgaccaagtgacagcaatgctgtttcactggttatgcggcggatccgaaaagaaaacgttgatgccggtgaacgtgcaaaacaggctctagcgttcgaacgcactgatttcgaccaggttcgttcactcatggaaaatagcgatcgctgccaggatatacgtaatctggcatttctggggattgcttataacaccctgttacgtatagccgaaattgccaggatcagggttaaagatatctcacgtactgacggtgggagaatgttaatccatattggcagaacgaaaacgctggttagcaccgcaggtgtagagaaggcacttagcctgggggtaactaaactggtcgagcgatggatttccgtttctggtgtagctgatgatccgaataactacctgttttgccgggtcagaaaaaatggtgttgccgcgccatctgccaccagccagctatcaactcgcgccctggaagggatttttgaagcaactcatcgattgatttacggcgctaaggatgactctggtcagagatacctggcctggtctggacacagtgcccgtgtcggagccgcgcgagatatggcccgcgctggagtttcaataccggagatcatgcaagctggtggctggaccaatgtaaatattgtcatgaactatatccgtaacctggatagtgaaacaggggcaatggtgcgcctgctggaagatggcgattaggcttatatgaagatgaagatgaaatatttggtgtgtcaaataaaaagcttgtgtgcttaagtttgtgtttttttcttggcttgttgtgttatgaatttgtggctttttctaatattaaatgaatgtaagatctcattataatgaataaacaaatgtttctataatccattgtgaatgttttgttggatctcttctgcagcatataactactgtatgtgctatggtatggactatggaatatgattaaagataag
