## Supplementary File 2 for "A Heat-Shock Inducible System for Flexible Gene Expression in Cereals"

**Supplementary Figures and Tables**

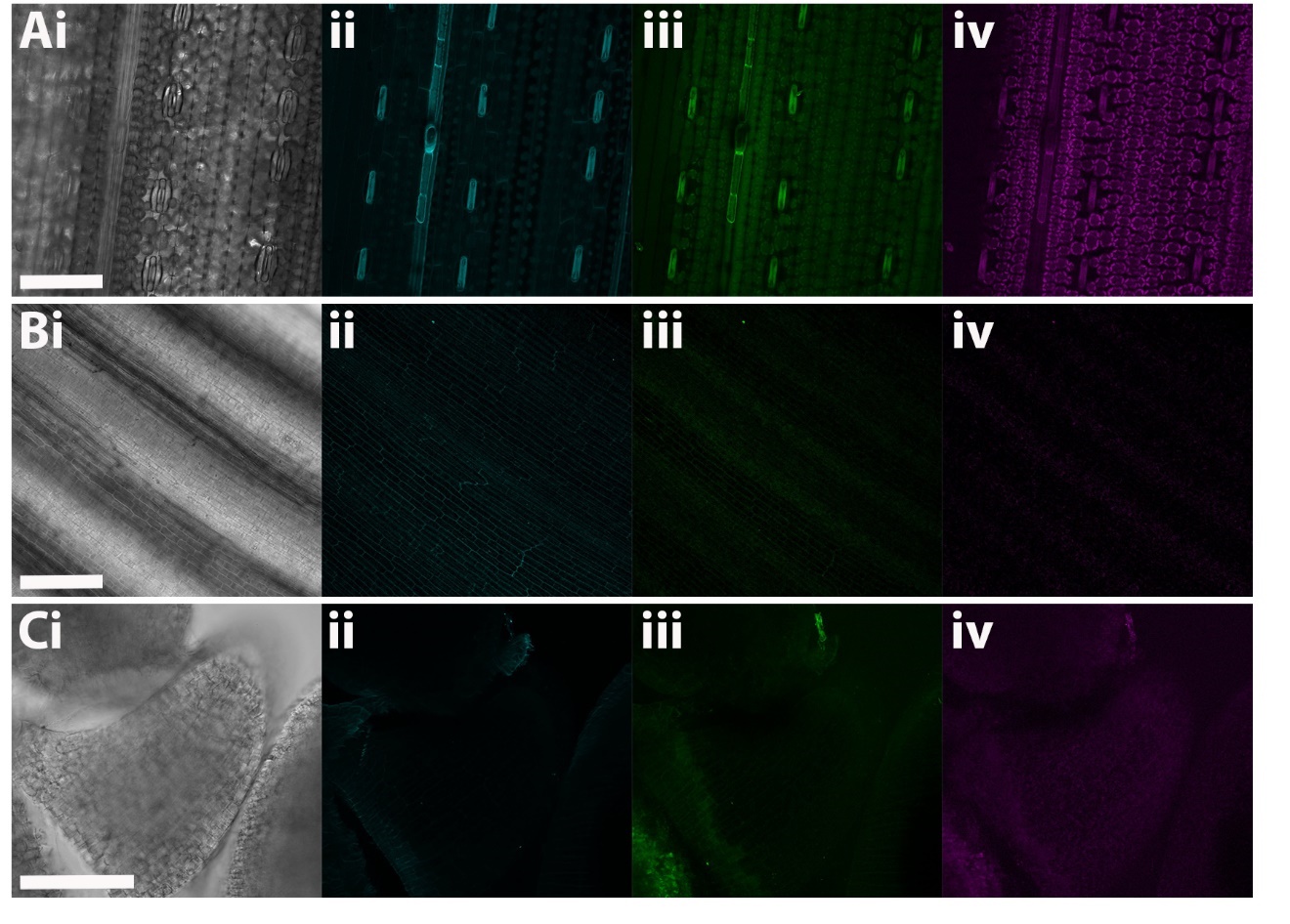

**Supplementary Figure 1: Screening of wild-type barley tissue for endogenous fluorescence.** *A-C.i:* bright field channels. *ii:* CyPET channel. *iii:* eGFP channel. *iv:* mCHERRY channel. *A.i-iv*: images of mature barley leaf tissue (blade); *B i-iv*: images of young barley leaf tissue; *C i-iv:* images of a young wild-type barley lemma. Gain > 120 for each tissue. Scale bars are 100µm.

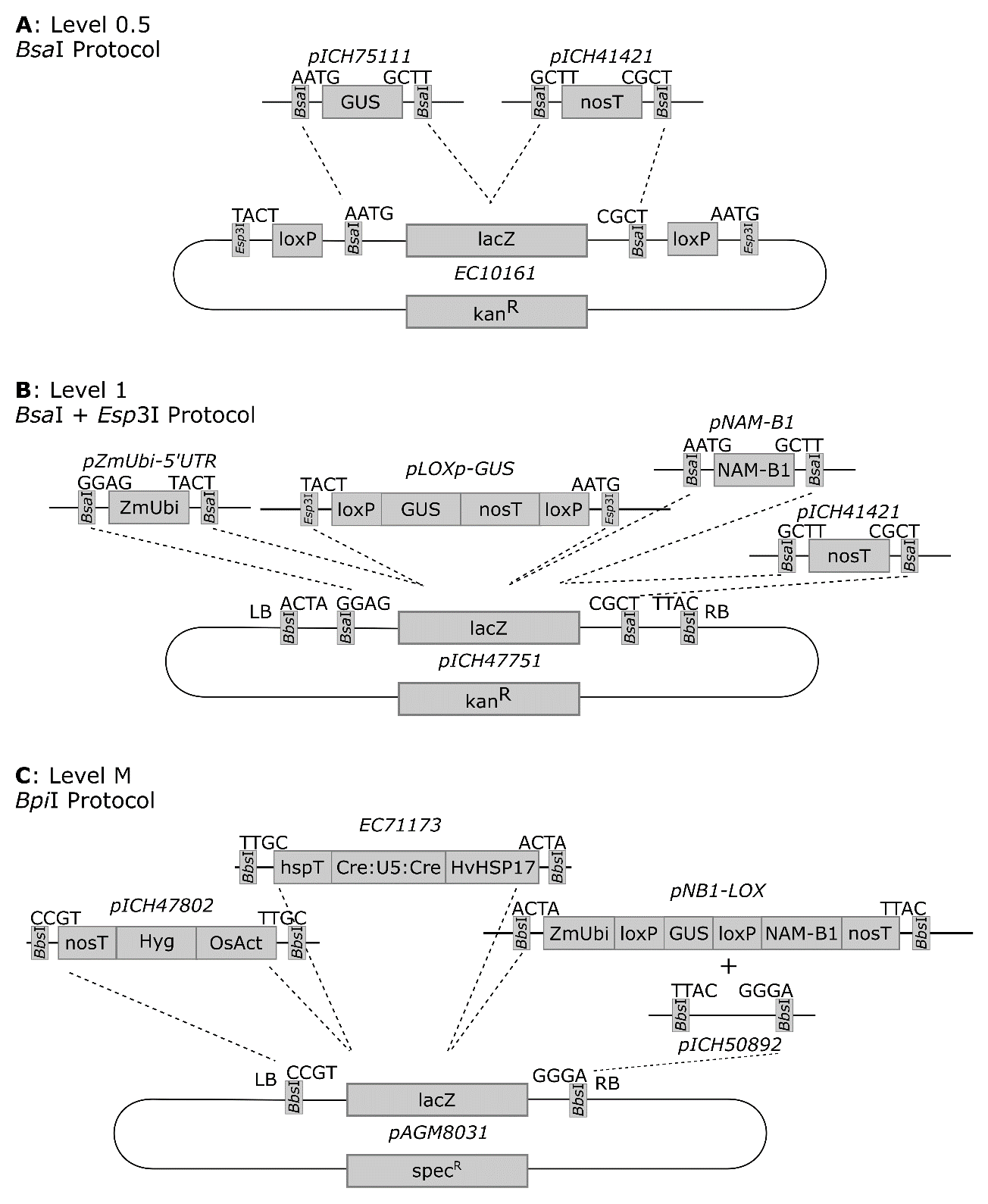

**Supplementary Figure 2: Construction of the wheat construct.** A) The reporter gene GUS (pICH75111) and the *nos* terminator (pICH41421) were cloned into the Level 0 vector EC10161, using the Type IIS restriction enzyme *Bsa*I. The resulting Level 0.5 vector contains loxP sites flanking the inserted reporter gene and terminator. B) The maize Ubiquitin promoter *ZmUbi* with a 5’ UTR sequence (from pICSL12009, cloned into pUAP1), *lox*P-flanked GUS (pLOXp-GUS), domesticated *NAM-*B1 in the universal acceptor pUAP1 (pNAM-B1), and the *nos* terminator (pICH41421) are cloned into a Level 1, position 3 vector (pAGM8031) in a reaction involving both *Bsa*I, for the canonical Level 0 and Level 1 parts, and *Esp*3I, for the *lox*P-containing Level 0.5 vector (see A).C) The hygromycin selection cassette (pICH47802), the HvHSP17::Cre-U5-Cre construct (EC71173), and the pNB1-LOX construct are cloned into a Level M, position 1 construct (pAGM8031), along with the position 4 end linker (pICH50892), using the Type IIS restriction enzyme *Bpi*I.

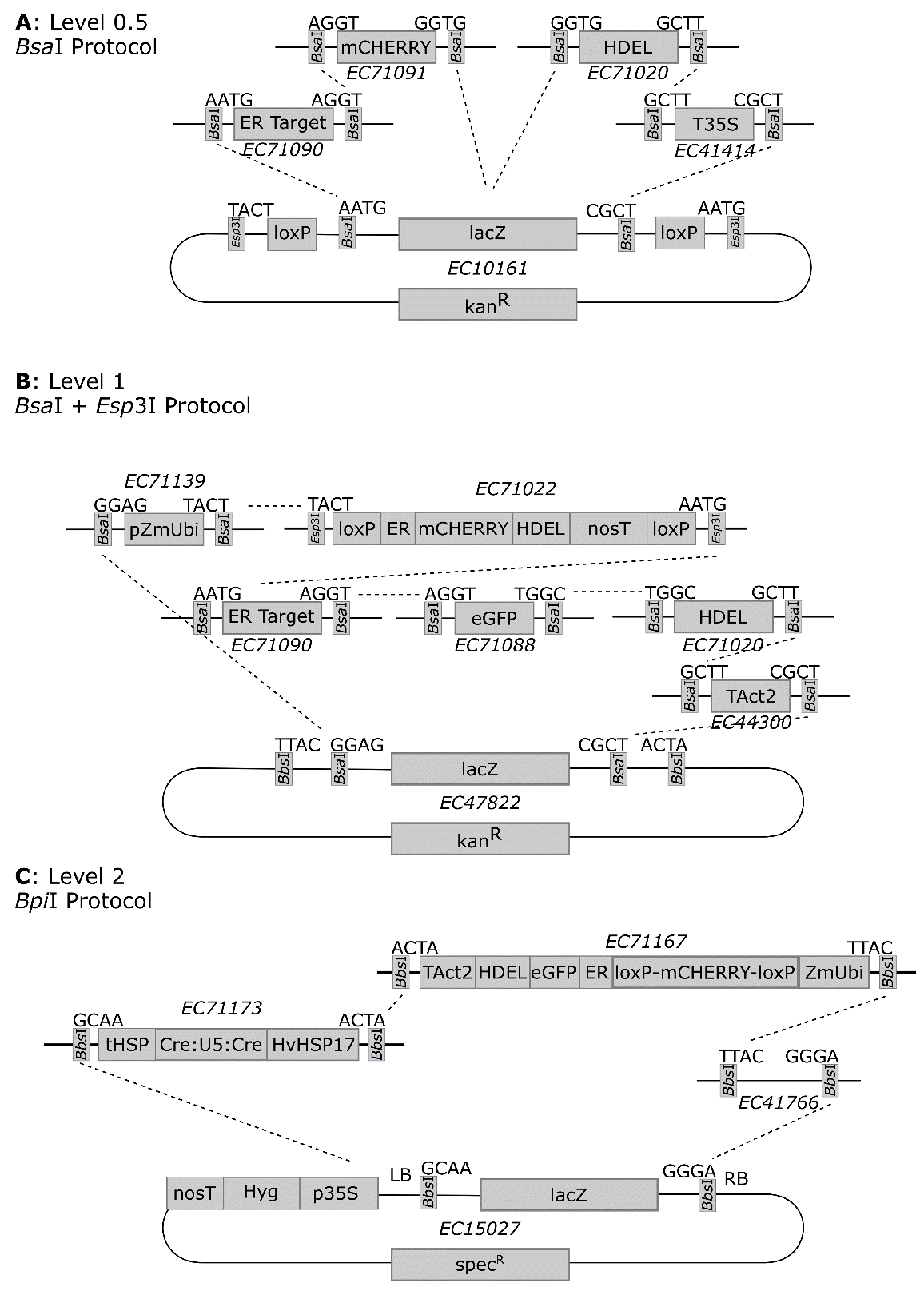

**Supplementary Figure 3: Construction of the barley construct.** A) The reporter gene mCHERRY (EC71091), an ER-targeting sequence (EC71090), a HDEL ER-targeting sequence (EC71020), and the 35S terminator (EC41414) were cloned into the Level 0 vector EC10161, using the Type IIS restriction enzyme *Bsa*I. The resulting Level 0.5 vector contains loxP sites flanking the inserted reporter gene and terminator. B) The maize Ubiquitin promoter *ZmUbi* with a 5’ UTR sequence (EC71139), *lox*P-flanked mCHERRY (EC10161), the ER-targeting sequence (EC71090), eGFP (EC71088), a HDEL ER-targeting sequence (EC71020), and the Actin2 terminator (EC41414) are cloned into a Level 1, position 3 vector (EC47822) in a reaction involving both *Bsa*I, for the canonical Level 0 and Level 1 parts, and *Esp*3I, for the *lox*P-containing Level 0.5 vector (see A).C) The HvHSP17::Cre-U5-Cre construct (EC71173), and the pGFP-LOX construct (EC71167), along with the position 4 end linker (EC41766), are cloned into a Level 2 construct (EC15027) which contains the 35s::Hygromycin selection cassette in position 1 using the Type IIS restriction enzyme *Bpi*I.

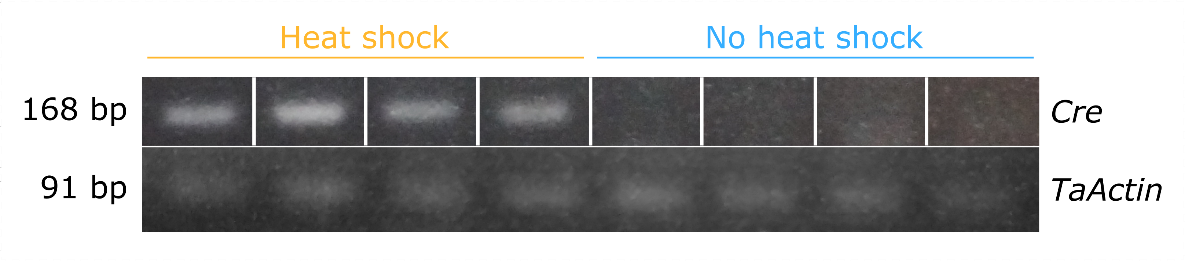

**Supplementary Figure 4: *Cre* expression in HS_NAM-B1 transgenic lines.** RT-PCR was carried out on each sample using primers SH033 and SH002 to amplify *Cre* and primers PB47 and PB48 to amplify the housekeeping gene *TaActin*; see primer sequences in Supplementary Table 3. The expected band size was 168 bp for *Cre* and 91 bp for *TaActin*. Bands showing *Cre* expression (top) were taken from the same gel and reordered for this figure and are therefore indicated as separate by the white vertical lines.

**Supplementary Table 1: Copy number of T_0_ transformants.** Lines taken forward for expression analysis and microscopy are highlighted in blue.

| **Construct** | **T_0_ Plant** | **Copies of Hygromycin** |
| --- | --- | --- |
| HS_NAM-B1 | 2020-20-01 | 46 |
|  | 2020-4-1 | 12 |
|  | 2020-19-01 | 10 |
|  | 2020-5-1 | 10 |
|  | 2020-13-1 | 4 |
|  | 2020-2-1 | 4 |
|  | 2020-8-1 | 4 |
|  | 2020-22-01 | 2 |
|  | 2020-23-01 | 2 |
|  | 2020-3-1 | 2 |
|  | 2020-21-01 | 1 |
|  | 2020-20-02 | 1 |
|  | 2020-24-01 | 1 |
|  | 2020-11-1 | 1 |
| HS_GFP | 00899-03 | 3-4 |
|  | 00899-02 | 2 |
|  | 00899-01 | 1 |
|  | 00899-04 | 1 |

**Supplementary Table 2: Constructs used in this study.** All constructs are listed with their name as used in the text, and in relevant databases, as well as a short description. The level of the construct, corresponding to those described in Weber et al. 2011 (Levels 0-2) and Werner et al. 2012 (Level M), is also provided.

| **Name** | **Description** | **Origin** | **Construct** | **Level** |
| --- | --- | --- | --- | --- |
| pICH41421 | NosT | The Sainsbury Lab (TSL) Synbio^1^ | Wheat | 0 |
| pICSL12009 | ZmUbi promoter + ubiquitin 5’ UTR (untranslated 1^st^ exon + intron) | TSL Synbio^1^ | Wheat | 0 |
| pICH75111 | GUS (β-glucuronidase gene) with 2 introns | TSL Synbio^1^ | Wheat | 0 |
| pNAM-B1 | Domesticated NAM-B1, with 2 introns | This study | Wheat | 0 |
| pZmUbi-5'UTR | ZmUbi promoter + ubiquitin 5’ UTR (untranslated 1st exon + intron), with L0.5-compatible overhangs | This study | Wheat | 0 |
| pICH47751 | Level 1 Position 3 | TSL Synbio^1^ | Wheat | 1 |
| pICH47802 | OsAct promoter + Hygromycin coding sequence, L1P1 | TSL Synbio^1^ and BRACT^2^ (Rey et al. 2018) | Wheat | 1 |
| pNB1-LOX | ZmUbi::loxP-GUS-nosT-loxP-NAM-B1-nosT; L1P3 | This study | Wheat | 1 |
| pAGM8031 | Level M Position 1 | TSL Synbio^1^ | Wheat | M |
| pICH50892 | Level M, end linker 3 | TSL Synbio^1^ | Wheat | MEL |
| pLOXp-GUS | loxP-GUS-nosT-loxP | This study | Wheat | 0.5 |
| HS_NAM-B1 | Level M, P1. OsAct::Hyg-nosT/HvHSP17::Cre-hspT/ZmUbi::loxP-GUS-loxP-NAM-B1-nosT |  |  |  |
| EC10161 | LoxP Vector | This study | Wheat and Barley | 0 |
| EC71173 | Level 1 P2 HspHv17::Cre-U5-Cre | This study | Wheat and Barley | 1 |
| EC47822 | L1 vector backbone pL1V-R3-47822 | ENSA^3^ | Barley | 1 |
| EC71139 | P-pZmUBI-intron L0 | This study | Barley | 0 |
| EC71022 | U-LoxP-mCHERRY-HDEL-t35S-loxP. L0.5 | This study | Barley | 0.5 |
| EC71090 | pL0M-S-ER-Targ_71090 | ENSA^3^ | Barley | 0 |
| EC71088 | pL0M-C1-eGFP-71088 | ENSA^3^ | Barley | 0 |
| EC71020 | pL0M-C2-HDEL-71020 HDEL | ENSA^3^ | Barley | 0 |
| EC44300 | T-Act2 | ENSA^3^ | Barley | 0 |
| EC41414 | T-35S | ENSA^3^ | Barley | 0 |
| EC71091 | pL0M-C1-mCherry | ENSA^3^ | Barley | 0 |
| EC71102 | Unmodified CRE L0. pL0M-SC-Cre | This study | Barley | 0 |
| EC71167 | pL1M-R3-UbqP-Loxp-ER-Targ-mCherry-HDEL-Loxp-eGFP | This study | Barley | 1 |
| EC71174 | pL2M-HvHSP17-CREU5-UBQ-loxmCHERRYER-eGFPER | This study | Barley | 2 |
| EC47811 | L1 vector backbone pL1V-R2-47811 | ENSA^3^ | Barley | 1 |
| EC71100 | pL0M-PU-pHvHSP17 | This study | Barley | 0 |
| EC71171 | CRE with U5 intron at 254bp, in pICH41308 backbone | This study | Barley | 0 |
| EC15320 | T-AtHsp | ENSA^3^ | Barley | 0 |
| pICSL80006 | Turbo GFP with intron from the arabidopsis U5 small nuclear ribonucleoprotein component gene | TSL Synbio^1^ | Barley | 0 |
| pICH41308 | Vector backbone for new CRE with intron | TSL Synbio^1^ | Barley | 0 |
| EC15027 | Level 2 Vector, including Hygromycin cassette at position 1 | ENSA^3^ | Barley | 2 |
| EC41766 | Level 2, end linker 3, pL1M-ele-3-41766 | ENSA^3^ | Barley | 2EL |

^1^https://www.synbio.tsl.ac.uk

^2^<https://www.jic.ac.uk/research-impact/technology-platforms/genomic-services/crop-transformation/>

^3^https://www.ensa.ac.uk/resources (Note that ENSA has no facility for directly distributing constructs, however ENSA will put interested researchers in contact with those who can distribute the constructs.)

**Supplementary Table 3: Primers used in the course of this study.** Calculated primer efficiencies for qRT-PCR are given where appropriate. For Level 0 cloning primers, the enzyme recognition sites are highlighted in blue, the resulting 4 base-pair overhangs are highlighted in red, and, where applicable, the Level 1 fusion sites are highlighted in bold.

| **Primer** | **Description** | **Sequence (5′ - 3′)** | **Reference** | **Efficiency** |
| --- | --- | --- | --- | --- |
| *gDNA_F* | Construct Excision PCR | CGATGCTCACCCTGTTGTTT | This study | NA |
| *gDNA_R* | Construct Excision PCR | GGTGATACCGCGTTGCTTTT | Borrill, P. (2013) | NA |
| *TaActin_F* | Actin, forward | ACCTTCAGTTGCCCAGCAAT | Uauy et al. (2006b) | 102% |
| *TaActin_R* | Actin, reverse | CAGAGTCGAGCACAATACCAGTTG | Uauy et al. (2006b) |  |
| *NAM-B1_F* | Domesticated NAM-B1, forward | AGTTGAACGGGGTCGACGAT | This study | 89% |
| *NAM-B1_R* | Domesticated NAM-B1, reverse | CTGCTGCCTCTCTCAGGTTG | This study |  |
| *Cre_F* | Cre recombinase, forward | ACCGGCATCAACGTTTTCTT | This study | NA |
| *Cre_R* | Cre recombinase, reverse | AAATGCTCCTGTCCGTTTGC | This study | NA |
| NAMB1_Level0_Forward_AATG | Level 0 cloning of NAM-B1 | ACGAAGACATCTCA**AATG**GGCAGCTCCGACTCATC | This study | NA |
| NAMB1_Level0_Reverse_GCTT | Level 0 cloning of NAM-B1 | ACGAAGACATCTCG**AAGC**TCAGGGATTCCAGTTCACGC | This study | NA |
| ZmUbi_Level0_Forward_GGAG | Level 0 cloning of ZmUbi | ATGAAGACATCTCA**GGAG**GTGCAGCGTGACCCGGTCGT | This study | NA |
| ZmUbi_Level0_Reverse_TACT | Level 0 cloning of ZmUbi | ACGAAGACATCTCG**AGTA**CCTGCAGAAGTAACACCAAA | This study | NA |
| Cre_E1_Level0_Forward_AATG | Level 0 cloning of Cre-U5-Cre | TGAAGACCAAATGTCCAATTTACTGACCGTACAC | This study | NA |
| Cre_E1_Level0_Reverse_AGGT | Level 0 cloning of Cre-U5-Cre | CGAAGACTTACCTGAAGATATAGAAGATAATCGC | This study | NA |
| Cre_I1_Level0_Forward_AGGT | Level 0 cloning of Cre-U5-Cre | CGAAGACTCAGGTAAGTTTCTGCTTCTACCTTTGATA | This study | NA |
| Cre_I1_Level0_Reverse_GCAG | Level 0 cloning of Cre-U5-Cre | AGAAGACGCCTGCACATCAACAAATTTTGGTCAT | This study | NA |
| Cre_E2_Level0_Forward_GCAG | Level 0 cloning of Cre-U5-Cre | AGAAGACGTGCAGGCGCGCGGTCTGGCAGTAA | This study | NA |
| Cre_E2_Level0_Reverse_GCTT | Level 0 cloning of Cre-U5-Cre | GGAAGACCAAAGCCTAATCGCCATCTTCCAGCAG | This study | NA |
